# NRT2.2 is essential for nitrate uptake and carbon-nitrogen balance in *Nannochloropsis oceanica*

**DOI:** 10.64898/2026.11.02.763162

**Authors:** Anna Santin, Laura Verdesca, Alessandra Bellan, Miles Minio, Caterina Gerotto, Tomas Morosinotto, Giorgio Perin, Alessandro Alboresi

**Affiliations:** Department of Biology, University of Padova, 35131, Padova, Italy; Department of Life and Environmental Sciences, Marche Polytechnic University, 60131 Ancona, Italy

**Keywords:** Nitrate transporters, nitrogen starvation, carbon-nitrogen balance, lipid accumulation, marine microalgae, *Nannochloropsis oceanica*, NRT2

## Abstract

Nitrate is the dominant nitrogen source for marine microalgae, yet how it is taken up remains poorly understood in many ecologically and biotechnologically relevant species. Here we show that NITRATE TRANSPORTER 2.2 (NRT2.2) is the principal nitrate transporter of the oleaginous marine microalga *Nannochloropsis oceanica*. *N. oceanica* encodes only two NRT2 proteins and, unlike land plants and green algae, lacks a recognizable homolog of NITRATE ASSIMILATION-RELATED 2 (NAR2), the partner protein that green-lineage NRT2s require for activity and plasma membrane targeting. CRISPR knock-outs and confocal imaging showed that NRT2.2 is located at the plasma membrane and is required for nitrate uptake, whereas its paralog NRT2.1 cannot compensate for its loss. When nitrate was resupplied after nitrogen starvation, *nrt2.2* mutants did not take up nitrate and did not resume growth, while they recovered normally with ammonium. Their photosystem II efficiency fell below that of fully nitrogen-starved wild-type cells, and chlorophyll content was reduced, while total protein remained stable. Fixed carbon was redirected into storage, with lipids and carbohydrates accumulating above the levels of all strains after 3 days of nitrogen starvation. Loss of NRT2.2 therefore mimics a prolonged nitrogen starvation, even though nitrate is available. Because nitrate was present but could not be used, recovery from starvation depends on the capacity to take up nitrate, not only on its availability. In the ocean, where nitrate is depleted for long periods and resupplied episodically, NRT2.2 is thus likely to set how quickly *Nannochloropsis* can use new nitrate inputs and resume growth.

## INTRODUCTION

Nitrogen (N) is an essential macronutrient for all photosynthetic organisms, required for the synthesis of proteins, nucleic acids, chlorophylls and numerous metabolic cofactors. Its availability therefore directly determines growth and productivity, as its assimilation is tightly coordinated with carbon fixation and energy metabolism (Simionato et al., 2013; Santin et al., 2026). While the molecular networks controlling N uptake and signaling are well established in land plants (Wang et al., 2018; Maghiaoui et al., 2020; Cassan et al., 2023), the mechanisms of N transport in marine microalgae remain poorly understood (McCarthy et al., 2017; Busseni et al., 2019; Smith et al., 2019).

Marine phytoplankton can use a variety of inorganic and organic N sources, including nitrate (NO_3_^-^), ammonium (NH_4_^+^), nitrite and, in several taxa, urea and amino acids. The relative contribution of these sources depends on the species, their environmental availability and the energetic cost of their assimilation (Glibert, 2016). Although NH_4_^+^ is energetically cheaper to assimilate, its concentration in seawater is very low, from ∼0.3 µM in the euphotic zone to ∼0.01 µM below it, whereas NO_3_^-^, although often below 1 µM in surface water, is the largest pool of bioavailable N in the ocean (Gruber, 2008; Sanz-Luque et al., 2015). In both cases, concentrations are lower and more variable than in soil, where they can reach the millimolar range (Kabala et al., 2017). Marine microalgae must therefore acquire NO_3_^-^ efficiently even at very low concentrations, which makes membrane transport a critical step in N acquisition, storage and assimilation.

NO_3_^-^ transport across biological membranes relies on proton-coupled active transporters of the major facilitator superfamily (MFS), which are predicted to contain twelve transmembrane helices (von Wittgenstein et al., 2014; Rogato et al., 2015). Within the MFS, NO_3_^-^ transport is primarily mediated by members of two evolutionarily distinct families: the low-affinity NITRATE TRANSPORTER 1/PEPTIDE TRANSPORTER FAMILY (NRT1/NPF) and the high-affinity NITRATE TRANSPORTER 2 (NRT2) family. The two families differ in their transport properties and physiological roles. NRT1/NPF proteins generally mediate NO_3_^-^ uptake at higher external concentrations and have diversified to transport a broad range of substrates, including peptides, hormones and secondary metabolites in addition to NO_3_^-^ (Fan et al., 2017; Santin et al., 2021; Santin et al., 2024). NRT2 proteins, in contrast, are primarily associated with high-affinity NO_3_^-^ uptake in plants and green algae (Fernandez and Galvan, 2008; Xu et al., 2024), making them particularly important when NO_3_^-^ is scarce. In land plants and in the freshwater alga *Chlamydomonas reinhardtii*, high-affinity NO_3_^-^ uptake by NRT2 requires the accessory protein NITRATE ASSIMILATION-RELATED 2 (NAR2, also known as NRT3) (Zhou et al., 2000; Rexach et al., 2002). In contrast, NAR2 homologs are absent from algae with secondary plastids (Rogato et al., 2015), suggesting that NRT2-mediated NO_3_^-^ transport operates differently in these lineages. However, NRT2 mutants have not yet been characterized in any secondary endosymbiont, so how NRT2 proteins work in these organisms is still unclear.

*Nannochloropsis* is a genus of marine microalgae with a secondary red plastid, widely studied for its high lipid productivity and as a model for marine photosynthesis and N metabolism (Simionato et al., 2013; Fattore et al., 2021). Hereafter, *Nannochloropsis* refers to the genus in the broad sense, including *Microchloropsis* (formerly *N. gaditana* and *N. salina*). In *Nannochloropsis*, N deprivation induces extensive metabolic and structural changes, including reorganization of the photosynthetic apparatus and the accumulation of storage lipids, highlighting the central role of N availability in controlling cellular physiology (Simionato et al., 2013; Fattore et al., 2021). Genomic and transcriptomic analyses have identified candidate genes for NO_3_^-^ assimilation in *Nannochloropsis*, including NRT2 transporters (Radakovits et al., 2012), but their function and subcellular localization are unknown, so it is still unclear how NO_3_^-^ enters the cell.

Here, we characterized the two NRT2 transporters of *N. oceanica* and showed that NRT2.2 is located at the plasma membrane and mediates NO_3_^-^ uptake from the medium. Loss of NRT2.2 prevents cells from resuming growth on NO_3_^-^ after N starvation, establishing a direct link between NRT2-dependent NO_3_^-^ uptake and recovery from N limitation. Cells lacking NRT2.2 behave as if N-starved even though NO_3_^-^ is available: they lose chlorophyll and photosynthetic efficiency and store the fixed C as lipids and carbohydrates.

## MATERIALS AND METHODS

### Identification of *Nannochloropsis oceanica* NRT2 sequences

*N. oceanica* NRT2 sequences were identified by tBLASTn searches in NanDeSyn (https://nandesyn.single-cell.cn/), using the six NRT2 proteins of *Phaeodactylum tricornutum* as queries against the *N. oceanica* IMET1 genome (Wang et al., 2014); redundant hits were then removed. The retrieved sequences were analyzed with TMHMM 2.0 (https://services.healthtech.dtu.dk/services/TMHMM-2.0/) to verify the presence of the 10-12 transmembrane helices (TMHs) typical of NRT2 transporters. Genomic and coding sequences were confirmed by PCR amplification and sequencing (Table S1 and Data Set S1). The number of NRT2 genes was further verified by BLASTp against all gene models and tBLASTn against the genome assembly of *N. oceanica* CCMP1779 (JGI PhycoCosm, https://phycocosm.jgi.doe.gov/), which retrieved only *NRT2.1* and *NRT2.2*.

To search for NAR2/NRT3 partner proteins, *Arabidopsis thaliana* (AtNAR2.1 ID AT5G50200.1 and AtNAR2.2 ID AT4G24730.1) and *Chlamydomonas reinhardtii* (Cre09.g410900) proteins were used as queries for BLASTp against all available *Nannochloropsis* proteomes. NRT2.1 and NRT2.2, retrieved at 100% identity, served as positive controls of the proteome search.

### Phylogenetic analysis of NRT2 proteins

NRT2 homologs were retrieved by tBLASTn searches with default parameters and an E-value threshold of 1 × 10^-5^, using the NRT2 amino acid sequences of *N. oceanica* and diatoms as queries, against NCBI (https://blast.ncbi.nlm.nih.gov/), the JGI Genome Portal (https://genome.jgi.doe.gov/portal/), Ensembl Protists (https://protists.ensembl.org/index.html) and PLAZA Diatoms 1.0 (https://bioinformatics.psb.ugent.be/plaza/versions/plaza_diatoms_01/). To cover all major eukaryotic lineages, this dataset was complemented with NRT2 sequences from UniProt (prasinophytes, trebouxiophytes, Chlorophyceae, red algae, haptophytes and cryptophytes) and from the JGI PhycoCosm genomes of *Microchloropsis gaditana* CCMP1894 and *Vischeria* sp. C74, the closest relatives of *Nannochloropsis* with sequenced genomes. The *Tribonema minus* sequence A0A835YPG2 was excluded because of a probable annotation error (an internal tandem duplication of about 30 residues). The final dataset included 94 NRT2 sequences (Supplementary Data Set S2).

Sequences were aligned with MAFFT v.7 using the L-INS-i algorithm (Katoh and Standley, 2013), and gap-rich, poorly aligned positions were removed with ClipKIT in gappy mode (gap threshold 0.4; (Steenwyk et al., 2020), yielding 469 amino acid positions, 435 of which were parsimony-informative. A maximum likelihood (ML) tree was inferred with IQ-TREE v.2.0.7 (Minh et al., 2020) under the LG+F+R7 model, selected as the best-fitting model by ModelFinder according to the Bayesian information criterion (Kalyaanamoorthy et al., 2017). Branch support was assessed with 1000 ultrafast bootstrap replicates (Hoang et al., 2018) and 1000 replicates of the SH-like approximate likelihood ratio test (SH-aLRT; Guindon *et al*., 2010). The tree was rooted at the midpoint. Alternative hypotheses on the origin of the two *N. oceanica* NRT2s (a duplication specific to *Nannochloropsis versus* a duplication before the divergence of *Nannochloropsis* and *Microchloropsis*) were compared with the approximately unbiased (AU) test (Shimodaira, 2002) implemented in IQ-TREE.

### Structural prediction of *N. oceanica* NRT2s

As no experimental structure of an NRT2 transporter is available, structural models of *N. oceanica* NRT2s were built with Phyre v.2.2 (Powell et al., 2025), using as template the crystal structure of the major facilitator superfamily (MFS) glycerol-3-phosphate transporter GlpT of *Escherichia coli* (SCOP domain: d1pw4a; PDB: 1PW4). Models were visualized with UCSF ChimeraX (Goddard et al., 2018). To identify residues putatively involved in NO_3−_ binding, the *N. oceanica* NRT2s were aligned with *Aspergillus nidulans* NrtA (formerly CrnA), whose key residues have been characterized by mutagenesis (Unkles et al., 2004), using the MUSCLE algorithm (Edgar, 2004) implemented in MEGA v.11.0.11 (Tamura et al., 2021).

### *N. oceanica* strain and culture

An axenic culture of *Nannochloropsis oceanica* strain IMET1 was obtained from the Provasoli-Guillard National Center for Marine Algae and Microbiota (www.bigelow.org). Cultures were maintained in F/2 medium (Guillard, 1975), prepared with 0.22 μm-filtered and autoclaved seawater, at 21 °C and 100 rpm, under continuous white fluorescent light (90 μmol photons m^−2^ s^−1^). Light intensity was measured with a LI-250A light meter (LI-COR, Lincoln, NE, USA).

### Generation and screening of *N. oceanica nrt2* knock-out and NRT2.2-mVenus lines

Guide RNAs (gRNAs) targeting *N. oceanica NRT2.1* and *NRT2.2* for Cpf1 (Cas12a) genome editing were designed with CRISPOR (http://crispor.tefor.net/crispor.py) on the *N. oceanica* IMET1 reference genome. Three crRNAs were selected for each gene, targeting regions predicted to encode TMHs to increase the likelihood of disrupting NRT2 function (the complete list of gRNAs is given in Table S1). The gRNAs were purchased from IDT (https://eu.idtdna.com/pages) and co-transformed with a linearized antibiotic resistance cassette, following the protocol of Perin *et al*. (2015). For selection, the *Aph7* gene from *Streptomyces hygroscopicus* (kindly provided by Prof. Krishna K. Niyogi (Vieler et al., 2012), conferring resistance to hygromycin B, was used under the control of the endogenous LDSP_E (Lipid Droplet Surface Protein E) promoter and the 35S terminator (Berthold et al., 2002; Vieler et al., 2012).

To generate NRT2.2-mVenus lines, *N. oceanica* IMET1 was co-transformed by electroporation with two cassettes: one for hygromycin resistance (Berthold et al., 2002; Vieler et al., 2012) required for selection, and one for NRT2.2-mVenus expression. The NRT2.2-mVenus plasmid was generated by Gibson Assembly (Gibson et al., 2009). In this plasmid, the NRT2.2 coding sequence was fused at its 3’ end to mVenus, followed by a His-tag, under the control of the endogenous *LDSP_E* promoter (Berthold et al., 2002; Vieler et al., 2012). Cloning primers are listed in Table S1. Both cassettes were excised from their plasmids and purified on a 0.8% (w/v) agarose gel. Transformation followed the protocol of Perin *et al*. (2015). Resistant colonies were picked after 4-5 weeks and screened by PCR with the primers listed in Table S1.

### Subcellular localization of NRT2.2

Cells were imaged with a Zeiss LSM 700 confocal laser scanning microscope (Carl Zeiss, Jena, Germany) equipped with a Plan-Apochromat 63x/1.40 Oil DIC M27 objective. mVenus was excited at 488 nm and detected at 524-536 nm, and chlorophyll *a* autofluorescence was excited at 640 nm and detected at 655-667 nm. Laser power, detector gain and offset were identical for all acquisitions. Single optical sections were acquired with ZEN Blue v.3.14 software (Carl Zeiss) at 1024 × 1024 pixels (0.099 µm pixel^-1^; WT and line #11) or 512 × 512 pixels (0.198 µm pixel^-1^; line #9).

Fluorescence distribution was quantified on the original 16-bit images with custom R scripts (image quantification) and Python scripts (profile analysis and figure assembly). In each image, the median background was subtracted from both channels and intensities were divided by the pixel dwell time (counts µs^-1^), so that images acquired with different settings could be compared. Bleed-through of chlorophyll *a* autofluorescence into the mVenus channel was estimated on WT cells, which do not express mVenus, as a constant fraction of the chlorophyll signal (4.1%), and subtracted from the mVenus channel of all images. In total, 32 cells from 15 independent fields were analyzed (WT, n = 10; NRT2.2-mVenus #9, n = 9; NRT2.2-mVenus #11, n = 13). In cells showing a peripheral mVenus ring, the cell center and ring diameter were determined by fitting a circle to the ring. Intensity profiles of mVenus and chlorophyll *a* were extracted along 18 diameters through the cell center and averaged for each cell. Peripheral enrichment was calculated as the ratio between the mean mVenus intensity within ±0.3 µm of the ring radius and that within the inner half of the radius. Co-localization of mVenus with chlorophyll *a* was assessed with Pearson’s correlation coefficient calculated on the bleed-through-corrected images.

### Growth under different N sources and concentrations

*N. oceanica* wild-type (WT) and knock-out (KO) lines were maintained in autoclaved filtered F/2 medium (Guillard, 1975) enriched in NH_4_^+^ (and 882 µM NH_4_Cl) in addition to the standard addition of 882 µM NaNO_3_. After three days (mid-exponential phase), cells were transferred to N-free F/2 medium for 72 h to deplete residual N. After 72 h of N starvation, cells were diluted to 1 × 10^7^ cells mL^-1^ in F/2 medium supplemented with 882 μM NaNO_3_, 250 μM NaNO_3_ or 882 μM NH_4_Cl. The experiment was performed first in 20 mL Erlenmeyer flasks and then in 80 mL tubes in a Multi-Cultivator MC 1000-OD (Photon Systems Instruments, Drásov, Czech Republic), under atmospheric CO_2_ and continuous white light (35 and 80 μmol photons m^-2^ s^-1^, respectively). Each condition was tested in at least three biological replicates.

Growth was monitored as optical density at 750 nm (OD_750_) with a Spark multimode microplate reader (Tecan, Männedorf, Switzerland) and confirmed by cell counting with an automated cell counter (Cellometer Auto X4, Nexcelom Bioscience, Lawrence, MA, USA). In the Multi-Cultivator, OD_680_ was recorded automatically every hour.

To measure extracellular NO_3_^−^ consumption, 1 mL of culture was filtered by vacuum filtration onto glass microfiber Whatman GF/F filters with nominal pore size 0.7 µm (Sigma-Aldrich, St. Louis, MO, USA), and the absorbance of the filtrate was measured at 220 nm with a Cary 60 UV-Vis spectrophotometer (Agilent Technologies, Santa Clara, CA, USA). NO_3_^−^ concentrations were calculated from a standard curve (10 μM to 1 mM NO_3_^−^ in N-free F/2 medium), according to McCarthy *et al*. (2017).

Data are presented as mean ± SD of independent biological replicates. Differences between each KO line and the WT were tested with an unpaired two-tailed Student’s t-test (for growth and NO_3_^−^ uptake, at day 4). Statistical analyses were performed with Python (SciPy).

### Pigments and protein extraction and quantification

Chlorophylls *a* (Chl *a*) and total carotenoids (Car) were extracted in the dark from culture samples with 100% N,N-dimethylformamide (DMF) (Sigma-Aldrich, St. Louis, MO, USA), following Liistro *et al*. (2026). Pigments in the extracts were quantified with an Agilent Cary 60 spectrophotometer (Agilent Technologies, Santa Clara, CA, US) using specific extinction coefficients (Moran, 1982). Absorbance at 750 nm was subtracted from all readings to correct for light scattering, and pigment content was expressed per cell using the cell concentration of each culture.

A total of 2 × 10^8^ cells were harvested from each culture, washed twice, and resuspended in 180 µL B1 buffer (20 mM Tris-KOH, pH 7.8, 400 mM NaCl, 2 mM MgCl_2_, 3 mM benzamidine, 1 mM PMSF). Cells were disrupted with acid-washed 150-212 µm glass beads (Sigma-Aldrich, St. Louis, MO, USA) using a Bullet Blender Storm Pro (Next Advance, Inc., Troy, NY, USA) for four cycles of 30 sec at 3500 rpm, with 3 min on ice between cycles. Subsequently, 80 µL of 3X sample buffer (45 mM Tris-HCl, pH 6.8, 30 mM DTT, 3% SDS, 10% glycerol) were added, followed by an additional 30 sec disruption step.

After extraction, proteins were precipitated with 4 volumes of cold 100% acetone for 1 h at -20 °C. Samples were centrifuged for 10 min at 15000 g at 4°C. The pellet was dried and then resuspended in 80 µl of B1 buffer (without protease inhibitors). Protein quantification was performed using a Bradford assay. For each sample, 33 µl were added to 1 ml of Bradford Reagent (Sigma-Aldrich, St. Louis, MO, USA) and incubated for 10 min at room temperature in the dark. Absorbance was measured at 595 nm. Protein concentrations were calculated from a bovine serum albumin (BSA) standard curve and expressed per cell, based on the 2 × 10^8^ cells extracted per sample.

### Determination of photosynthetic efficiency

Chlorophyll fluorescence was measured *in vivo* with a Dual-PAM-100 fluorometer (Heinz Walz, Effeltrich, Germany). After 25 min of dark adaptation, the maximum quantum yield of Photosystem II (PSII) was calculated as F_v_/F_m_ = (F_m_−F_0_)/F_m_ (Maxwell and Johnson, 2000), where F_0_ is the minimum fluorescence of dark-adapted cells and F_m_ the maximum fluorescence measured during a saturating light pulse (∼7000 μmol photons m^−2^ s^−1^, 600 ms).

### Quantification of intracellular neutral lipids

Intracellular neutral lipids were quantified by Nile Red staining (Sigma-Aldrich, St. Louis, MO, USA). 1 × 10^7^ cells were collected, resuspended in 200 µL of F/2 medium and then stained with 0.625 µg mL^−1^ Nile Red (DMSO stock solution) for 15 min at 37 °C in the dark with shaking (100 rpm). Fluorescence was measured with the Tecan microplate reader described above (excitation 485 nm, emission 580 nm) and corrected for background fluorescence using unstained samples and water blanks.

### Elemental C and N composition

For C and N quantification, cell pellets were washed three times with an isosmotic ammonium formate solution (0.5 M) and freeze-dried. Capsules containing 0.8-1.2 mg of dried sample were analyzed with a CHN elemental analyzer (ECS 4010, Costech, Italy) coupled to an ID Micro EA isotope ratio mass spectrometer (Compact Science Systems, Newcastle-under-Lyme, UK) to determine C and N content as a percentage of sample dry weight. Urea was used as the analytical standard.

In the *nrt2.2* KO lines grown with NO_3_^−^, N was below the detection limit of 4% N/DW in all but one replicate; as the single measurable value was itself above the detection limit, these samples were reported as having %N below this threshold (Fig. 3C). Because N could not be detected, the instrument software did not calculate %C for these samples. %C was therefore calculated from the C peak areas as follows, after verifying that this calculation reproduced the values automatically reported for the samples in which %C had been provided:

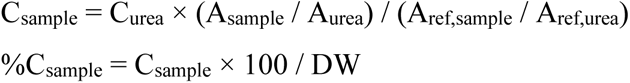

### Fourier-transform infrared (FTIR) spectroscopy

About 1 mg of dried cells, previously washed three times with the isosmotic ammonium formate solution, was weighed and resuspended in 110 μL of grade 1 water; 50 μL of the suspension were placed on a silicon disk and dried at 80 °C. FTIR spectra of whole cells were acquired in a range from 400 to 4000 cm^−1^, sampling every 1.93 cm^−1^ with a Tensor 27 FTIR spectrometer (Bruker Optics GmbH, Ettlingen, Germany). FTIR bands were assigned to cellular pools of proteins, carbohydrates, and lipids as described in Giordano *et al*. (2001). The band at about 1656 cm^-1^ was assigned to protein, the one at about 1740 cm^-1^ to lipids and the ones in the range 1160-1030 cm^-1^ to carbohydrates. Band integrals of the deconvolved spectra were obtained using the OPUS v.6.5 software (Bruker Optics GmbH, Ettlingen, Germany) and used to calculate the relative abundances of proteins, carbohydrates and lipids. For each strain and condition, three biological replicates were analyzed in technical duplicates, and technical duplicates were averaged before statistical analysis.

## RESULTS

### 1. Identification, evolution and structure of *N. oceanica* NRT2 transporters

tBLASTn searches against the *N. oceanica* IMET1 genome (Wang et al., 2014) identified two NRT2 sequences, NO30G01230 and NO14G02450, hereafter referred to as NRT2.1 and NRT2.2 respectively (Data Set S1). Both gene models were supported by available transcriptomic data and confirmed by PCR.

To place the *N. oceanica* NRT2 proteins in an evolutionary context, we reconstructed a maximum-likelihood phylogeny of 94 NRT2 proteins from stramenopiles, the green lineage, red algae, haptophytes, cryptophytes and fungi (Fig. 1A; Supplementary Data Set S2). The tree resolved two strongly supported groups: one comprised sequences from land plants and green algae together with those of haptophytes, red algae and cryptophytes, whereas the other included stramenopiles and fungi (Fig. 1A). In the tree, NRT2.2 groups with the single NRT2 of *Microchloropsis*, whereas NRT2.1 branches outside this pair (Fig. 1A). Together with an AU test rejecting a duplication within *Nannochloropsis* (p = 0.002), this suggests that the two *N. oceanica* NRT2s arose from a duplication that occurred before the divergence of *Nannochloropsis* and *Microchloropsis*, followed by the loss of NRT2.1 in *Microchloropsis*.

We generated structural models of *N. oceanica* NRT2.1 and NRT2.2 using Phyre2.2 (template: bacterial MFS transporter d1pw4a, PDB: 1PW4). Both models predicted 12 transmembrane helices (TMHs), with the N- and C-termini in the cytoplasm (Fig. 1B). Since the two models were highly similar, NRT2.2 is shown as representative (Fig. 1B), and the corresponding NRT2.1 residues are reported in Fig. S1. Mutagenesis studies of *Aspergillus nidulans* NrtA (previously CrnA) (Unkles et al., 2004) identified key residues required for NO_3_^-^ binding. Two positively charged arginine residues, R86 in TMH2 and R302 in TMH8, are conserved in NRT2.2 and correspond to residues required for NO_3_^-^ binding in NrtA (Fig. 1B; Fig. S1 for NRT2.1). Three further conserved residues were identified in the same putative binding site: N167 in TMH4, which may contribute to substrate stabilization; E265, whose negative charge may balance the positive charges of R86 and R302 through a salt bridge; and N390, likely involved in positioning NO_3_^-^ or maintaining the conformation of the translocation pathway (Fig. 1B) (Unkles et al., 2004). All five residues are also conserved in NRT2.1 (Fig. S1). The conservation of the complete binding-site architecture in both paralogs indicates that both proteins could, in principle, transport NO_3_^-^.

**Figure 1.**
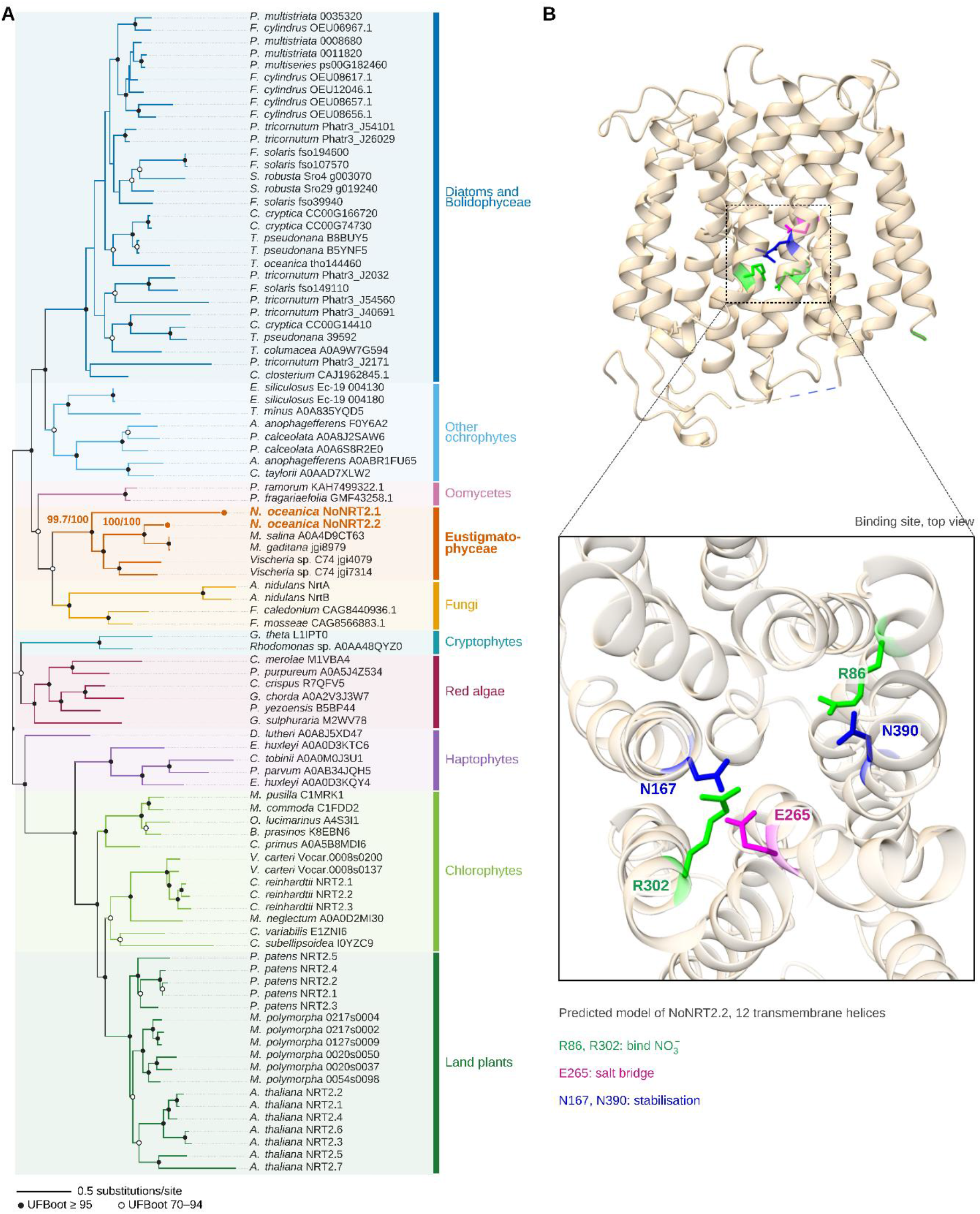
Phylogeny and predicted structure of *Nannochloropsis oceanica* NRT2s. A) Maximum-likelihood phylogeny of 94 NRT2 proteins from 50 photosynthetic and non-photosynthetic eukaryotes. Filled and open circles indicate nodes with ultrafast bootstrap (UFBoot) support ≥ 95% and 70-94%, respectively; SH-aLRT/UFBoot values are shown for the eustigmatophyte clade and for the NoNRT2.2-*Microchloropsis* clade. Colors indicate taxonomic groups; *N. oceanica* NRT2.1 and NRT2.2 are in orange. Sequences and accession numbers are listed in Supplementary Data Set S1. B) Predicted structural model of *N. oceanica* NRT2.2 with 12 transmembrane helices and, in the zoomed view, the putative NO_3_^-^ binding site: R86 and R302 (green), E265 (magenta), and N167 and N390 (blue). ALT TEXT: Phylogenetic tree of NRT2 transporters and predicted 3D structure of *Nannochloropsis* NRT2.2.

### 2. Loss of NRT2.2 abolishes NO_3_^-^ uptake in *N. oceanica*

To dissect the functional role of the two NRT2 transporters, we generated *N. oceanica nrt2.1* and *nrt2.2* knock-out (KO) mutants using a CRISPR-Cpf1 genome editing approach, targeting each gene with three different gRNAs (Fig. S2). Amplification and sequencing of the target loci confirmed the mutations (Fig. S2). In the *nrt2.1* KO lines #1 and #2, the hygromycin resistance cassette was integrated at the target locus. *nrt2.2* KO line #1 carries the cassette in the antisense orientation relative to NRT2.2, whereas line #2 carries the cassette in the sense orientation (Fig. S2). RT-PCR analysis confirmed the absence of detectable *NRT2.1* or *NRT2.2* transcripts in two independent KO lines per gene (Fig. 2A), which were selected for phenotypic characterization.

The KO lines were isolated and maintained in medium containing both NO_3_^-^ and NH_4_^+^, in which *nrt2.1* and *nrt2.2* KO strains grew similarly to the wild type (WT) (data not shown), as NH_4_^+^ can sustain growth independently of NO_3_^-^ transport. To test NO_3_^-^ utilization specifically, cells were starved of N for 3 days to deplete intracellular N reserves and then resupplied with either NO_3_^-^ or NH_4_^+^ as the sole N source (Fig. 2B). Upon resupply of 882 µM NO_3_^-^, extracellular NO_3_^-^ progressively decreased in WT and *nrt2.1* KO cultures, to <100 µM after 4 days, whereas it remained unchanged in both *nrt2.2* KO lines, demonstrating a complete loss of NO_3_^-^ uptake (Fig. 2C). In WT cultures ∼20% of the NO_3_^-^ was already taken up after 1 day, before any increase in cell number (Fig. 2C,D), indicating that the lack of NO_3_^-^ uptake in *nrt2.2* KO lines is not a consequence of impaired growth. Accordingly, WT and *nrt2.1* KO cultures resumed growth, tripling cell density within 4 days, whereas both *nrt2.2* KO lines showed no increase in cell number (Fig. 2D). The same phenotype was observed at a lower NO_3_^-^ concentration (250 µM): *nrt2.2* KO lines neither took up NO_3_^-^ nor grew, whereas WT and *nrt2.1* KO cells reduced extracellular NO_3_^-^ by ∼70% within 4 days (Fig. S3A,B).

In contrast, when NH_4_^+^ was resupplied as the sole N source, *nrt2.2* KO strains recovered similarly to WT and *nrt2.1* KO (Fig. 2E,F), indicating that their growth defect is specific to NO_3_^-^ and is not a general effect of the mutations. These findings identify NRT2.2 as an essential component of the NO_3_^-^ uptake machinery in *N. oceanica*, required for NO_3_^-^ acquisition and NO_3_^-^-dependent growth.

**Figure 2.**
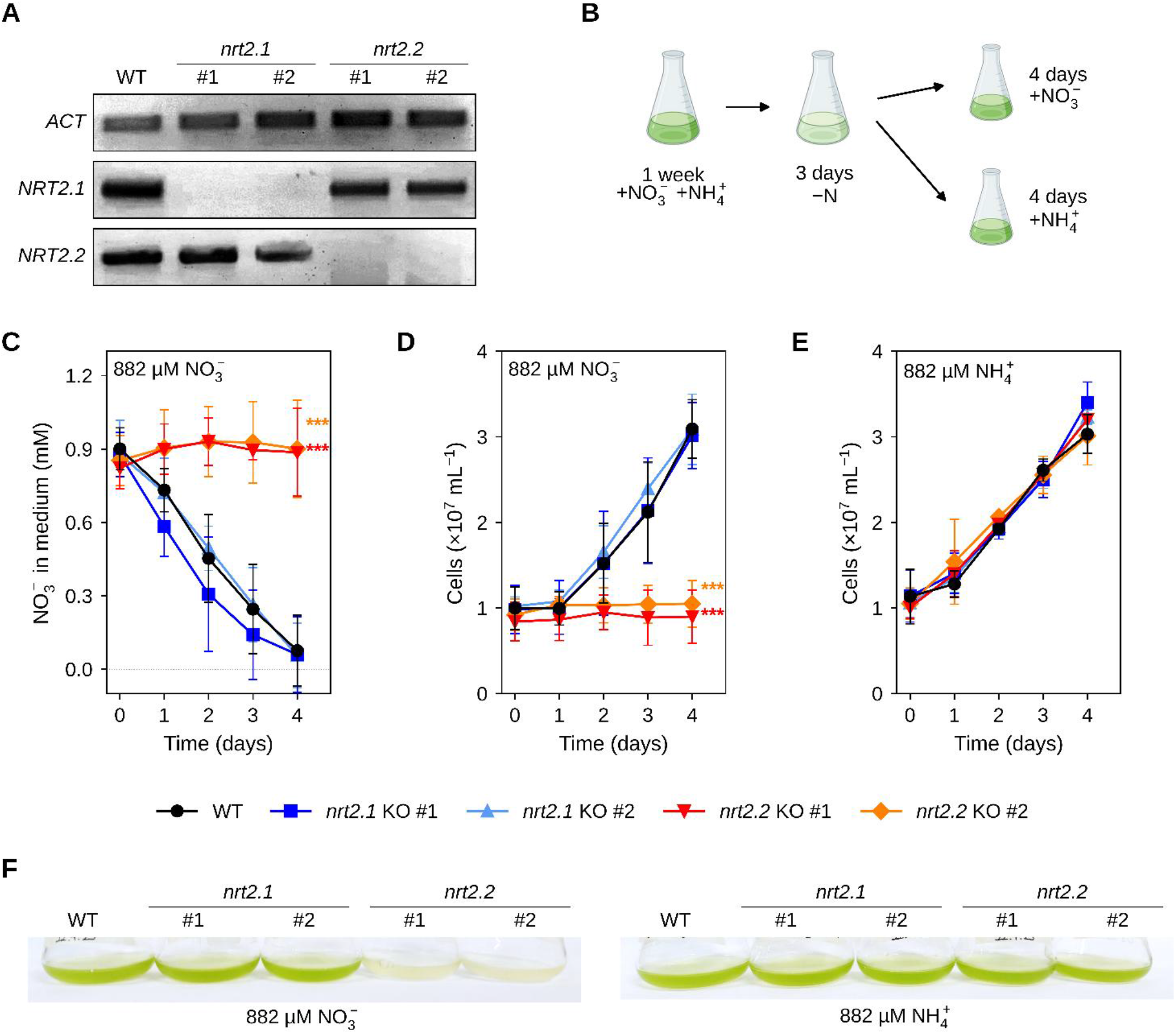
Impaired growth and nitrate uptake in *Nannochloropsis oceanica nrt2.2* KO mutants. A) RT-PCR of NRT2.1 and NRT2.2 transcripts in the wild type (WT) and in two *nrt2.1* and two *nrt2.2* knock-out (KO) lines (#1, #2); ACT (actin), reference transcript. Primers are listed in Table S1. B) Experimental setup. Cells were grown under atmospheric CO_2_ (400 ppm) and continuous white light (35 µmol photons m^-2^ s^-1^) in 882 µM nitrate (NO_3_^-^) and 882 µM ammonium (NH_4_^+^) until exponential phase, starved of N for 3 days and then resupplied with NO3-or NH_4_^+^. C) Residual NO_3_^-^ in the medium and D) cell concentration during 4 days in 882 µM NO_3_^-^. E) Cell concentration in 882 µM NH_4_^+^ (control). F) Cultures after 4 days of NO_3_^-^ or NH_4_^+^ repletion. Data are mean ± SD of at least three biological replicates. Asterisks: unpaired two-tailed Student’s t-test vs WT at day 4; * p < 0.05, ** p < 0.01, *** p < 0.001. ALT TEXT: Gel, scheme and growth curves showing that *nrt2*.2 knock-out lines do not take up nitrate and do not grow on it.

### 3. Intracellular C and N partitioning following loss of NRT2.2

To understand the consequences of the loss of NO_3_^-^ uptake on cell physiology, we characterized the C and N composition, storage metabolism and photosynthetic performance of WT, *nrt2.1* KO and *nrt2.2* KO strains. To obtain enough biomass, cells were grown in a Multi-Cultivator photobioreactor under the same starvation-repletion regime and sampled at the end of the 3-day N depletion (hereafter -N) and after 4 days of repletion with NH_4_^+^ or NO_3_^-^ (hereafter +NH_4_^+^ and +NO_3_^-^), the same time point used in flasks, when WT cultures had consumed most of the NO_3_^-^ (Fig. 2C). As in flasks, *nrt2.2* KO lines neither grew (Figs. 3A and S3C,D) nor consumed NO_3_^-^ (Fig. S4) in +NO_3_^-^, whereas all strains behaved like WT in -N and +NH_4_^+^ (Fig. S4). Since *nrt2.2* KO cells cannot use NO_3_^-^, at the end of the +NO_3_^-^ treatment they had been without an available N source for 7 days (3 days of depletion followed by 4 days of repletion), compared with 3 days for -N samples.

The lack of NO_3_^-^ uptake caused a strong intracellular N deficiency. In +NO_3_^-^, *nrt2.2* KO cells showed a slight increase in total carbon (C) content relative to WT and *nrt2.1* KO (Fig. 3B) and a strong reduction in N content, with five of six samples falling below the detection limit of the assay (Fig. 3C), yielding a sharply elevated C/N ratio (Fig. S5). Since most cellular N is allocated to proteins, we then quantified total protein content, which was unaffected across genotypes and conditions (Figs. 3D and S6). *nrt2.2* KO cells therefore maintained their protein content per cell while accumulating C. Since N is expressed per dry weight, its decrease largely reflects dilution by C-rich compounds. In parallel, maximum PSII efficiency (F_v_/F_m_) was strongly reduced in *nrt2.2* KO cells in +NO_3_^-^ (0.43-0.49 *vs* 0.65 in WT; Fig. 3E), well below the values of all strains in -N (∼0.61) and +NH_4_^+^ (Fig. S6), without an available N source (7 vs 3 days). Chlorophyll, a major N sink of the photosynthetic apparatus, was reduced by up to ∼50% in *nrt2.2* KO cells, to levels similar to −N (Figs. 3F and S7), and the overall pigment composition shifted (Fig. 3G). The decrease in F_v_/F_m_ in *nrt2.2* KO cells was thus accompanied by a loss of chlorophyll, while total protein per cell was maintained.

**Figure 3.**
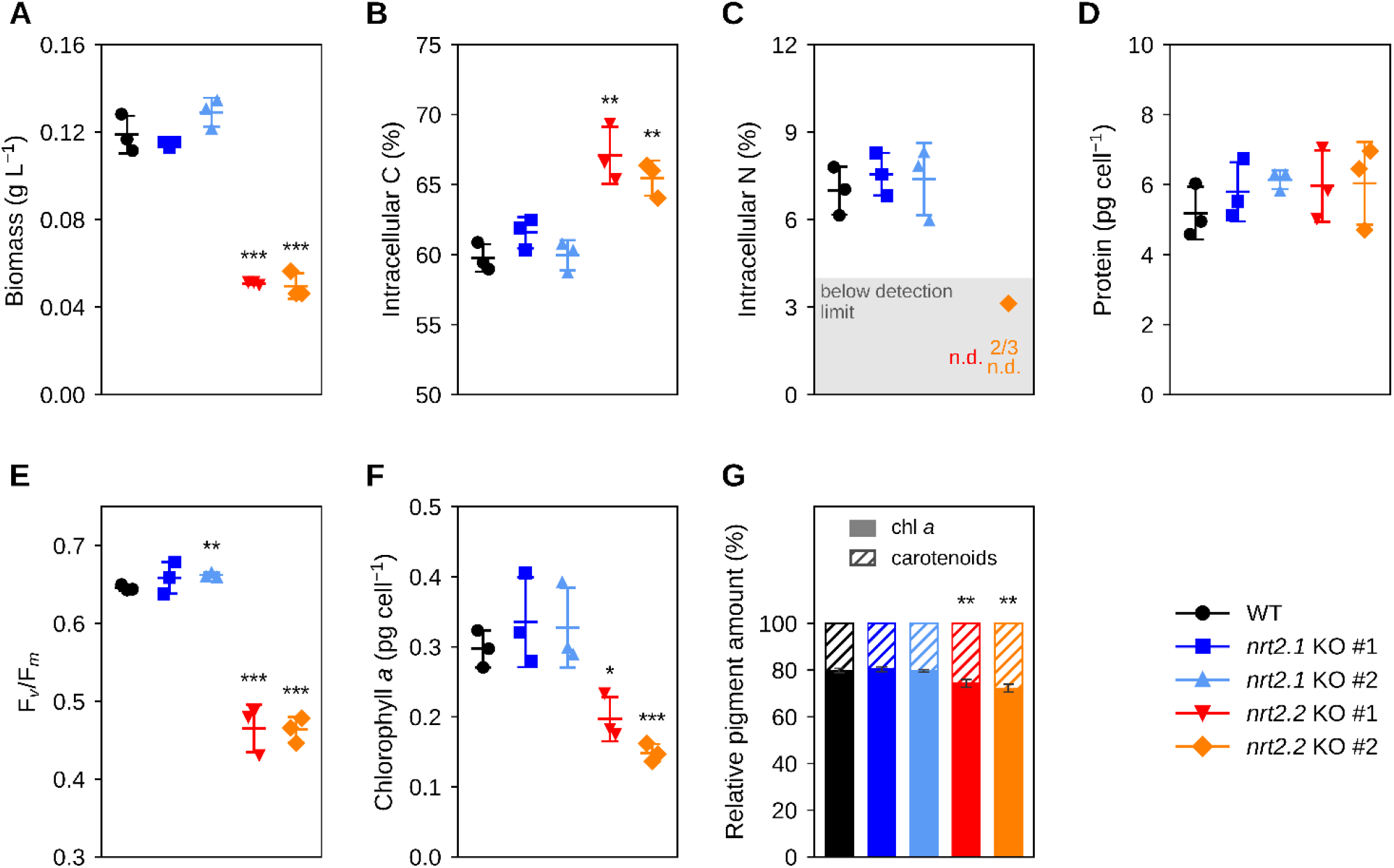
Altered intracellular N content and photosynthetic performance in *Nannochloropsis oceanica nrt2.2* knock-out mutants. Wild-type (WT), two *nrt2.1* and two *nrt2.2* knock-out (KO) lines were grown in the Multi-Cultivator system under atmospheric CO_2_ (400 ppm) and continuous white light (80 µmol photons m^-2^ s^-1^) and analyzed after 4 days of repletion with 882 µM nitrate (NO_3_-); growth curves are shown in Figure S3 and the control conditions, nitrogen (N) deprivation and NH_4_^+^ repletion, in Figures S4-S7. A) Biomass production. B) Carbon (C) and C) N content, as percentage of dry weight. The detection limit for N is 4% of dry weight (grey area): N was not detected (n.d.) in five of six *nrt2.2* KO samples, and the only measurable sample (*nrt2.2* KO #2) is shown. D) Total protein content. E) Maximum photosynthetic efficiency, expressed as F_v_/F_m_, and F) chlorophyll content. G) Relative pigment composition (% chlorophyll and total carotenoids). Data are mean ± SD of at least three biological replicates. Asterisks: unpaired two-tailed Student’s t-test vs WT; * p < 0.05, ** p < 0.01, *** p < 0.001. ALT TEXT: Graphs showing higher C/N and lower photosynthetic efficiency in *nrt2.2* knock-out lines.

We next asked where the excess C was stored. To answer this, whole cells were analyzed by Fourier transform infrared (FTIR) spectroscopy, which measures the absorption of the main functional groups of cellular macromolecules: the amide I band of proteins (∼1650 cm^-1^), the ester C=O band of lipids (∼1740 cm^-1^) and the C-O-C bands of carbohydrates (950-1200 cm^-1^) (Fig. 4A-C). Because total protein content was stable across strains and conditions (Figs. 3D and S6), spectra were normalized to the amide I band, and lipid/protein and carbohydrate/protein ratios were quantified by fitting the spectra with Lorentzian bands (Fig. 4D,E). After 3 days of N depletion, all strains accumulated lipids, with a lipid/protein ratio about twice that in +NH_4_^+^, while the carbohydrate/protein ratio did not increase (Fig. 4A,B,D,E). In both conditions, the ratios of *nrt2.2* KO lines were similar to WT; the only differences, in single *nrt2.1* KO lines, were not confirmed in the second line (Fig. 4D,E). In +NO_3_^-^, by contrast, *nrt2.2* KO spectra diverged from all other strains, with a clear increase in both bands (Fig. 4C). The lipid/protein ratio of *nrt2.2* KO lines was ∼3.7-fold higher than in WT (Fig. 4D), and the carbohydrate/protein ratio ∼1.4-fold higher (Fig. 4E).

Neutral lipids were also measured independently by Nile Red staining, a fluorescent dye that binds neutral lipids such as triacylglycerols in intact cells. In all strains, Nile Red fluorescence was ∼2.5-fold higher after N depletion than in +NH_4_^+^ (Fig. 4F). In +NO_3_^-^, *nrt2.2* KO lines showed ∼3-fold higher fluorescence than WT, reaching or slightly exceeding N-depleted levels (Fig. 4F). C fixation therefore continues in *nrt2.2* KO cells, but in the absence of NO_3_^-^ uptake fixed C cannot be incorporated into N-containing compounds and is redirected into lipid and carbohydrate storage: an N-starvation signature, although NO_3_^-^ is available in the medium.

**Figure 4.**
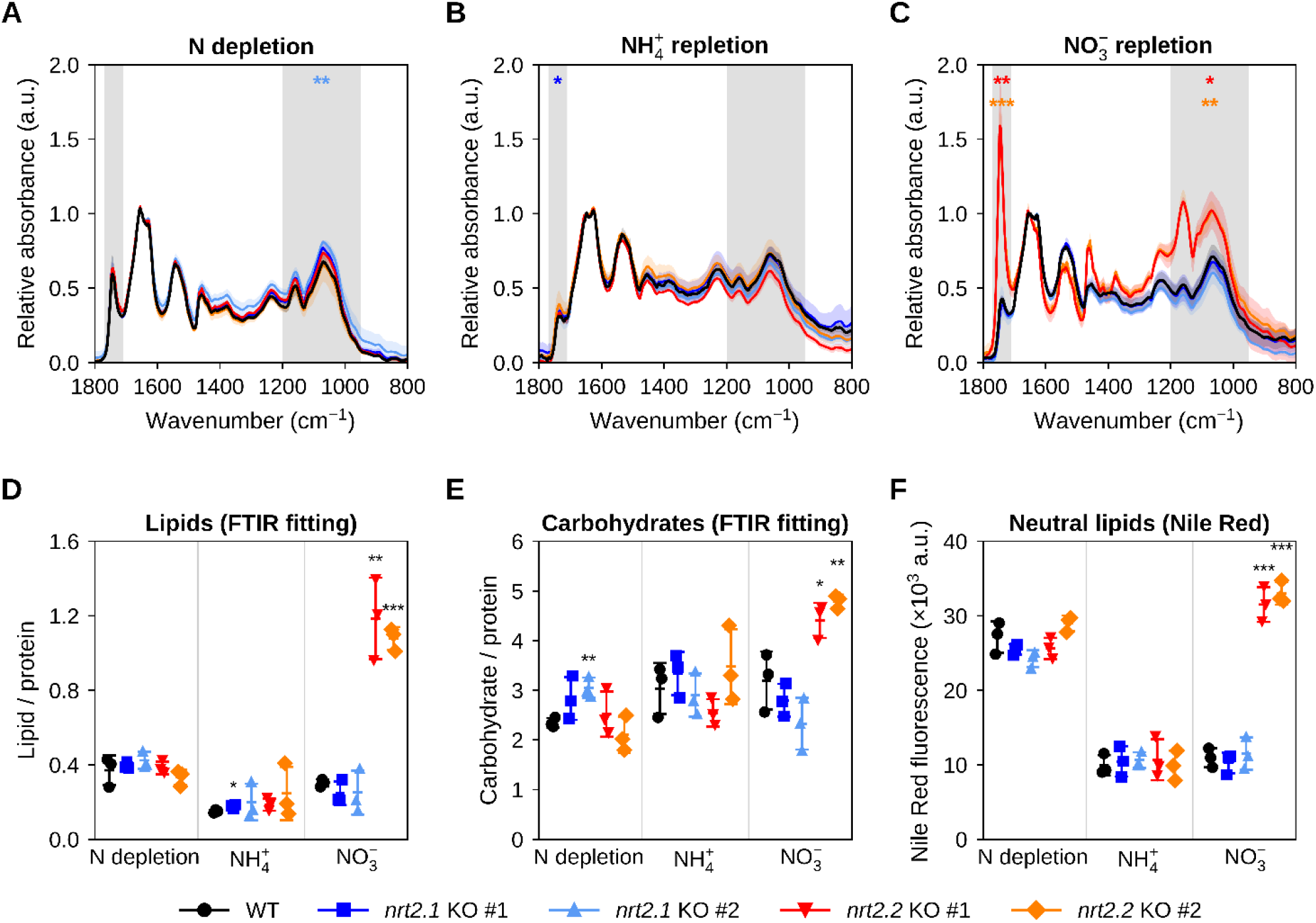
Increased lipid and carbohydrate content in *Nannochloropsis oceanica nrt2.2* knock-out mutants. Fourier Transform Infrared (FTIR) spectra of *N. oceanica* wild-type (WT), two *nrt2.1* and two *nrt2.2* knock-out (KO) lines grown in the Multi-Cultivator system, under atmospheric CO_2_ (400 ppm) and continuous white light (80 µmol photons m^-2^ s^-1^) A) after 3 days of nitrogen (N) depletion, B) after 4 days of repletion with 882 µM ammonium (NH4+), and C) after 4 days of repletion with 882 µM nitrate (NO_3_^-^). Spectra are normalized to the protein amide I peak at ∼1650 cm-1; total protein did not differ between strains (Fig. S6). Shaded areas mark the spectral regions of the fitted bands used for quantification: lipid ester C=O (1770-1710 cm^-1^) and carbohydrate C-O-C (1200-950 cm^-1^); colored asterisks refer to the band ratios in D and E. D) Lipid (ester C=O) and E) carbohydrate (C-O-C) band areas relative to the protein amide I band (∼1656 cm^-1^), obtained by fitting the spectra with Lorentzian bands (lipids: ester C=O band at ∼1740 cm^-1^; carbohydrates: sum of the bands between ∼1030 and ∼1160 cm^-1^). F) Neutral lipids, measured as Nile Red fluorescence, in the same conditions. Data are mean ± SD of at least three biological replicates (FTIR technical duplicates were averaged). Asterisks: unpaired two-tailed Student’s t-test vs WT; * p < 0.05, ** p < 0.01, *** p < 0.001. ALT TEXT: FTIR spectra and plots showing more lipids and carbohydrates in *nrt2.2* knock-out lines.

### 4. Subcellular localization of NRT2.2 on the plasma membrane

Since loss of NRT2.2 caused the strongest phenotype under NO_3_^-^ supply, we focused on this transporter and determined its subcellular localization. We generated stable lines expressing NRT2.2 fused at its C-terminus to mVenus (NRT2.2-mVenus) (Fig. 5A). After transformation, independent lines were screened by PCR for integration of the construct (Fig. 5B), and two positive lines (#9 and #11) were selected for confocal microscopy.

In both lines, mVenus formed a ring at the cell periphery, flanking the chlorophyll signal (Fig. 5C,D). mVenus fluorescence at the cell edge was 3.9 ± 1.2-fold (line #9, n = 4 cells) and 4.3 ± 0.9-fold (line #11, n = 5 cells) higher than in the cell interior, and the diameter of the fluorescent ring (3.1 ± 0.4 and 3.4 ± 0.5 µm, respectively) was consistent with the size of *N. oceanica* cells. In addition, the correlation between mVenus and chlorophyll signals within each cell was close to zero or slightly negative (mean Pearson’s r = −0.13 in both lines; Fig. 5E), indicating that the fusion protein is excluded from the chloroplast. Together, these data localize NRT2.2 to the plasma membrane, at the interface between the cell and the external medium, consistent with its role in NO_3_^-^ uptake from seawater in *N. oceanica*.

**Figure 5.**
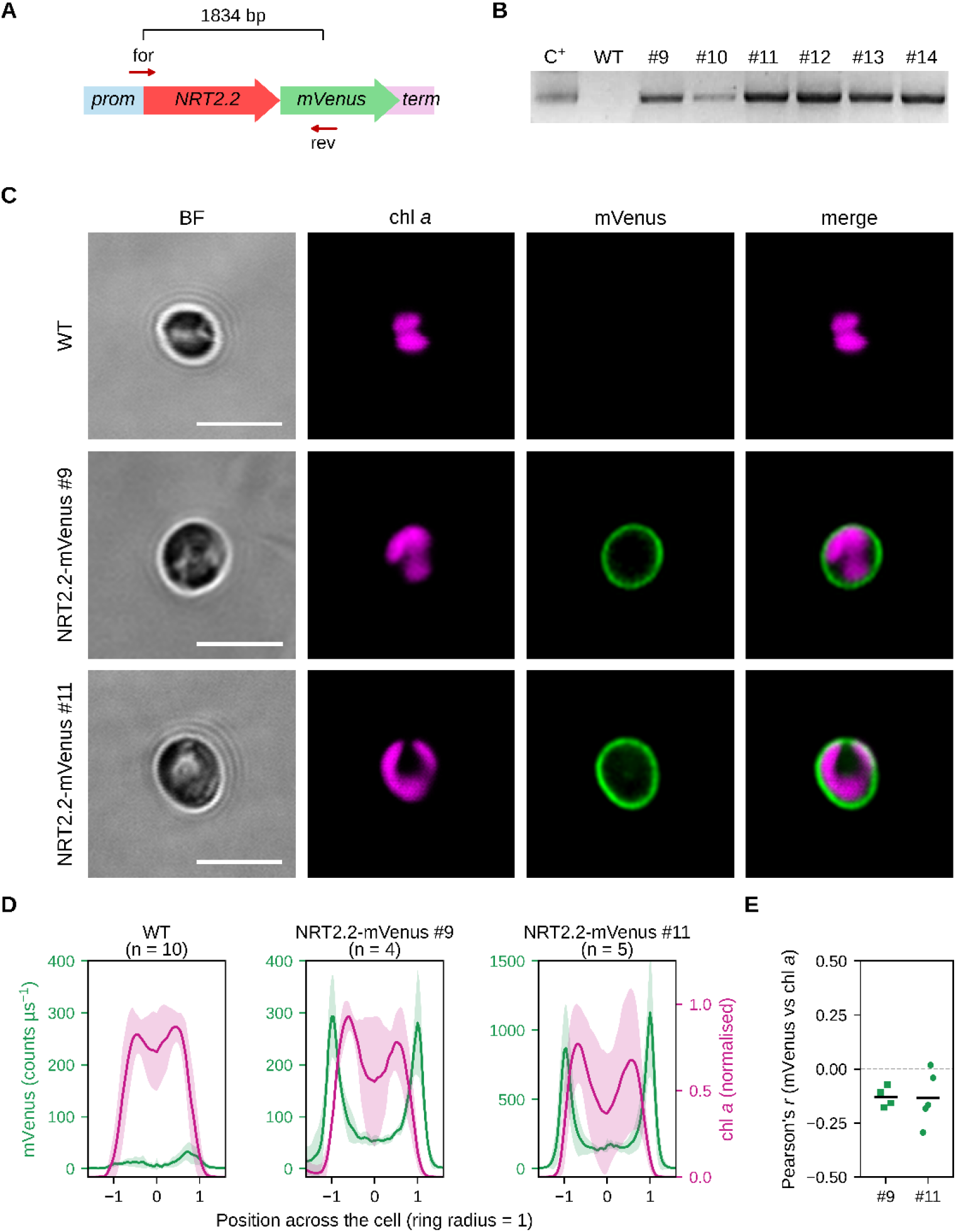
Subcellular localization of *Nannochloropsis oceanica* NRT2.2 to the plasma membrane. A) Expression construct: NRT2.2 coding sequence fused at its 3’ end to the yellow fluorescent protein mVenus, co-transformed with a hygromycin resistance cassette. B) Genomic PCR confirming the integration of the construct (1834 bp amplicon, primers for/rev in A) in lines #9 and #11; WT, wild-type parental strain; C+, transformation plasmid. C) Confocal images of WT and NRT2.2-mVenus cells: bright field (BF), chlorophyll *a* autofluorescence (magenta), mVenus (green) and merge. mVenus is corrected for chlorophyll bleed-through and shown with the same intensity range in all lines. Scale bars = 5 μm. D) Mean intensity profiles (± SD) of mVenus (green) and chlorophyll *a* (magenta, normalized to the maximum of each profile) across the cell (18 diameters per cell; position normalized to the ring radius, ring at ±1; WT scaled by the mean ring radius, 1.63 µm), in WT cells (n = 10) and in NRT2.2-mVenus cells with a peripheral ring (#9: n = 4; #11: n = 5); mVenus is on a separate scale for each line. E) Pearson’s correlation between mVenus and chlorophyll *a* signals after bleed-through correction in the NRT2.2-mVenus lines #9 and #11; each dot is one cell, bars are means. ALT TEXT: Confocal images showing NRT2.2-mVenus at the cell periphery.

## DISCUSSION

N availability in marine environments fluctuates in time and space, and N is often the primary limiting nutrient for marine primary production (Gruber, 2008; Sanz-Luque et al., 2015). Here we identified NRT2.2 as the plasma membrane transporter that mediates NO_3_^-^ uptake in *N. oceanica*. NRT2.2 is required for NO_3_^-^-dependent growth, and its loss disrupts the balance between C and N metabolism, driving cells into a physiological state that resembles prolonged N starvation, even though NO_3_^-^ is available.

### NRT2.2 is the main gateway for nitrate acquisition in *N. oceanica*

*N. oceanica* possesses only two *NRT2* genes, *NRT2.1* and *NRT2.2*, whereas the closely related *Microchloropsis* species carry a single one (Radakovits et al., 2012), although annotation inconsistencies in these genomes call for caution when estimating gene number (Fig. 1A). The single NRT2 retained in *Microchloropsis* is the ortholog of NRT2.2 (Fig. 1A), suggesting that the duplication occurred before the split between the two genera and was followed by the loss of NRT2.1 in *Microchloropsis*. This is consistent with NRT2.2 representing the core NO_3_^-^ transporter of the lineage, and NRT2.1 an accessory paralog. However, NRT2 sequences cluster mainly by taxonomic lineage (Rogato et al., 2015; Busseni et al., 2019), so phylogenetic position alone cannot predict the function of each paralog. This compact organization contrasts with that of land plants and green algae, in which NO_3_^-^ uptake relies on multiple, partially redundant NRT2 members with distinct regulatory profiles, such as AtNRT2.1 and AtNRT2.2, the major components of the high-affinity NO_3_^-^ transport system in *Arabidopsis thaliana* (Filleur et al., 2001; Li et al., 2007; Kotur et al., 2012), and the multiple, differentially regulated NRT2 homologs of *Chlamydomonas reinhardtii* (Fernandez and Galvan, 2008).

In land plants and *C. reinhardtii*, high-affinity NO_3_^-^ uptake by NRT2 requires the partner protein NAR2 (Zhou et al., 2000; Rexach et al., 2002), which is absent from *N. oceanica* and from other algae with secondary plastids (Rogato et al., 2015). The complete loss of NO_3_^-^ uptake in the *nrt2.2* KO lines (Fig. 2C) shows that NRT2.2 is functional *in vivo* without NAR2, either alone, like the fungal transporter NrtA, or with a partner protein not yet identified. However, whether NRT2.2 operates as a high-affinity transporter remains to be tested: the uptake assays used here were performed at 250-882 µM NO_3_^-^, well above typical seawater concentrations, and kinetic measurements at low micromolar NO_3_^-^ will be needed to determine its affinity. Beyond copy number, NRT2 proteins are conserved across eukaryotic lineages, and this conservation extends to the structural level. Although no eukaryotic NRT2 structure has been experimentally resolved, the predicted models of *N. oceanica* NRT2 showed the typical 12-TMH architecture and the conservation of charged and polar residues, suggesting that the molecular determinants required for NO_3_^-^ recognition and membrane transport remain highly conserved (Fig. 1B).

The knock-out mutants showed that one of the two copies has the main role: loss of NRT2.2 completely abolished NO_3_^-^ uptake (Fig. 2C) and growth on NO_3_^-^ (Fig. 2D), whereas growth on NH_4_^+^ was comparable to the WT (Fig. 2E). The defect is therefore specific to NO_3_^-^ acquisition and identifies NRT2.2 as the major gateway for extracellular NO_3_^-^ in this species. The absence of compensation by NRT2.1 indicates that, unlike in plants and green algae, NO_3_^-^ acquisition in *N. oceanica* relies on a single dominant transporter rather than on partially redundant family members. This specialization may reflect the ecological conditions of marine microalgae: unlike land plants, which face NO_3_^-^ gradients largely determined by soil properties and root architecture (Kabala et al., 2017), marine phytoplankton experience rapidly changing nutrient conditions, where NO ^-^ can shift from complete depletion in stratified, nutrient-poor waters to sudden enrichment after upwelling or mixing (Gruber, 2008; Sanz-Luque et al., 2015). In this setting, because NRT2.2 controls NO_3_^-^ entry, its capacity may determine how quickly *N. oceanica* resumes growth when NO_3_^-^ becomes available again.

No clear phenotype was found in the *nrt2.1* KO lines, and three explanations are possible. First, NRT2.2 may fully compensate for the loss of NRT2.1 (but not vice versa), as documented for NRT2 paralogs in vascular plants (Xu et al., 2024; Berman et al., 2026). Second, the two transporters may differ in affinity or kinetic properties (Liu et al., 1999). The NO_3_^-^ concentrations tested here (250-882 µM, much higher than in seawater), set by the resolution of the assay, may not have captured the activity range of NRT2.1. Third, the expression of *N. oceanica* NO_3_^-^ transporters responds to environmental conditions, including timing of N starvation (Li et al., 2014), light and C supply (Wei et al., 2019), so NRT2.1 may become relevant only under conditions not examined here.

Finally, the plasma membrane localization of NRT2.2 (Fig. 5) places it at the interface between the cell and its environment, consistent with its role in the uptake of extracellular NO_3_^-^.

### Loss of NRT2.2 disrupts the C/N balance and triggers a starvation-like metabolic state

The consequences of impaired NO_3_^-^ uptake extended beyond growth inhibition and highlighted a central role of NRT2.2 in maintaining cellular metabolic homeostasis. Disrupting the main NO_3_^-^ entry point produces an internal N deficit in cells that remain surrounded by NO_3_^-^, and the resulting phenotype can be read as the metabolic signature of N starvation, imposed here genetically rather than environmentally.

The most direct consequence of NRT2.2 disruption was the decrease of chlorophyll content and photosynthetic performance (Figs. 3F,G and 6), indicating that N limitation primarily affected the abundance and functional capacity of the photosynthetic apparatus. Chlorophyll is an N-rich molecule, and the light-harvesting complexes bind it in large amounts. Reducing pigment content could therefore represent a direct way to lower N demand without dismantling the protein scaffold of the photosystems (Wang et al., 2024). This is consistent with N deprivation triggering chloroplast remodeling, degradation of photosynthetic pigments and reduction of photosystem activity in microalgae (Bender et al., 2014; Fattore et al., 2021). In N-starved WT *Nannochloropsis*, this remodeling is an acclimation response: photosynthesis remains operational and supplies energy and reducing power for cell maintenance and lipid synthesis (Simionato et al., 2013; Wang et al., 2024). The *nrt2.2* KO lines instead face a permanent block of NO_3_^-^ uptake, and their phenotype reflects a chronic N deficiency that parallels, but does not necessarily reproduce, the response of WT cells to N starvation.

In *N. oceanica*, long-term N deprivation drives a global down-regulation of protein expression, with the induction of N-scavenging enzymes, proteases and proteasome components to recycle N from existing cellular components, and autophagy of plastids (Dong et al., 2013; Simionato et al., 2013). In the *nrt2.2* KO, by contrast, total protein content remained stable (Figs. 3D and 6): these lines reproduce chlorosis and reduced photosynthetic performance, but not the proteome-wide N recycling of N-starved cells, which in WT cells appears only after longer starvation. The sharp decrease in N content despite stable protein levels reflects the different normalization of the two measurements: N is expressed per dry weight and protein per cell, so the accumulation of C-rich lipids and carbohydrates (Fig. 4) increases cell dry weight and dilutes N, while non-protein N pools such as chlorophyll also decrease (Fig. 3F). Two non-exclusive interpretations can be proposed. First, N allocation may follow a hierarchy in which pigments are sacrificed before the protein pool, preserving the core cellular machinery. Second, the absence of bulk protein remobilization may indicate that *nrt2.2* KO cells do not fully engage the canonical N-recycling program of N-starved cells (Dong et al., 2013; Wang et al., 2024), so that the chronic deficit is not buffered by protein degradation. This makes the *nrt2.2* KO a useful tool to dissect which components of the starvation response depend on the perception of external N and which on the internal N deficit.

Another metabolic consequence of NRT2.2 loss was the accumulation of C-rich storage compounds, including neutral lipids and carbohydrates (Figs. 4 and 6). This matches the classical response to N deficiency and reflects the metabolic coupling between C and N. The reduction of NO_3_^-^ to NH_4_^+^ and its subsequent incorporation into amino acids through the GS-GOGAT cycle consume C skeletons (2-oxoglutarate), ATP and reducing power from photosynthesis. In the *nrt2.2* KO, NO_3_^-^ cannot enter the cell and none of these demands arises, while light-driven C fixation continues, albeit probably at a reduced capacity (Figs. 3C and 6) (Santin et al., 2026). The fixed C and reducing power that would normally feed amino acid synthesis are therefore redirected (McCarthy et al., 2017; Smith et al., 2019) toward storage compounds (Simionato et al., 2013; Fattore et al., 2021). This response is shared by the *nrt2.2* KO and by N-starved WT cells even though the two states differ in their pigment and protein responses, suggesting that C storage is a robust consequence of N deficiency irrespective of how the deficiency is imposed. Triacylglycerols (TAG) allow cells to store excess C in an osmotically neutral form (Goncalves et al., 2016).

Carbohydrates also increased in the *nrt2.2* KO lines (Fig. 4E). Carbohydrate accumulation is a common response to N limitation in microalgae (Pancha and Imamura, 2026). *Nannochloropsis*, however, stores C as β-1,3-glucan rather than starch (Vogler et al., 2018), and typically favors lipid over carbohydrate accumulation during N stress (Recht et al., 2012). The simultaneous increase of both pools in *nrt2.2* KO strains therefore suggests that the C surplus exceeded what the lipid pathway alone could absorb, or that the chronic *nrt2.2* block, with no demand for C and reducing power for NO_3_^-^ assimilation from the start, used both sinks. The two pools may also play distinct roles: carbohydrate synthesis is a rapid, low-cost sink for fixed C, whereas TAG synthesis requires more ATP and reducing power and represents a more reduced, long-term store (Valledor et al., 2014; Wang et al., 2024). Time-resolved measurements of both pools would clarify whether carbohydrates act as an early sink that is progressively complemented by lipids. Overall, the strong accumulation of both compounds indicates that the disruption of NO_3_^-^ uptake places cells in a prolonged C-rich, N-deficient metabolic state that cannot be relieved just by NO_3_^-^ supply (Fig. 2).

**Figure 6.**
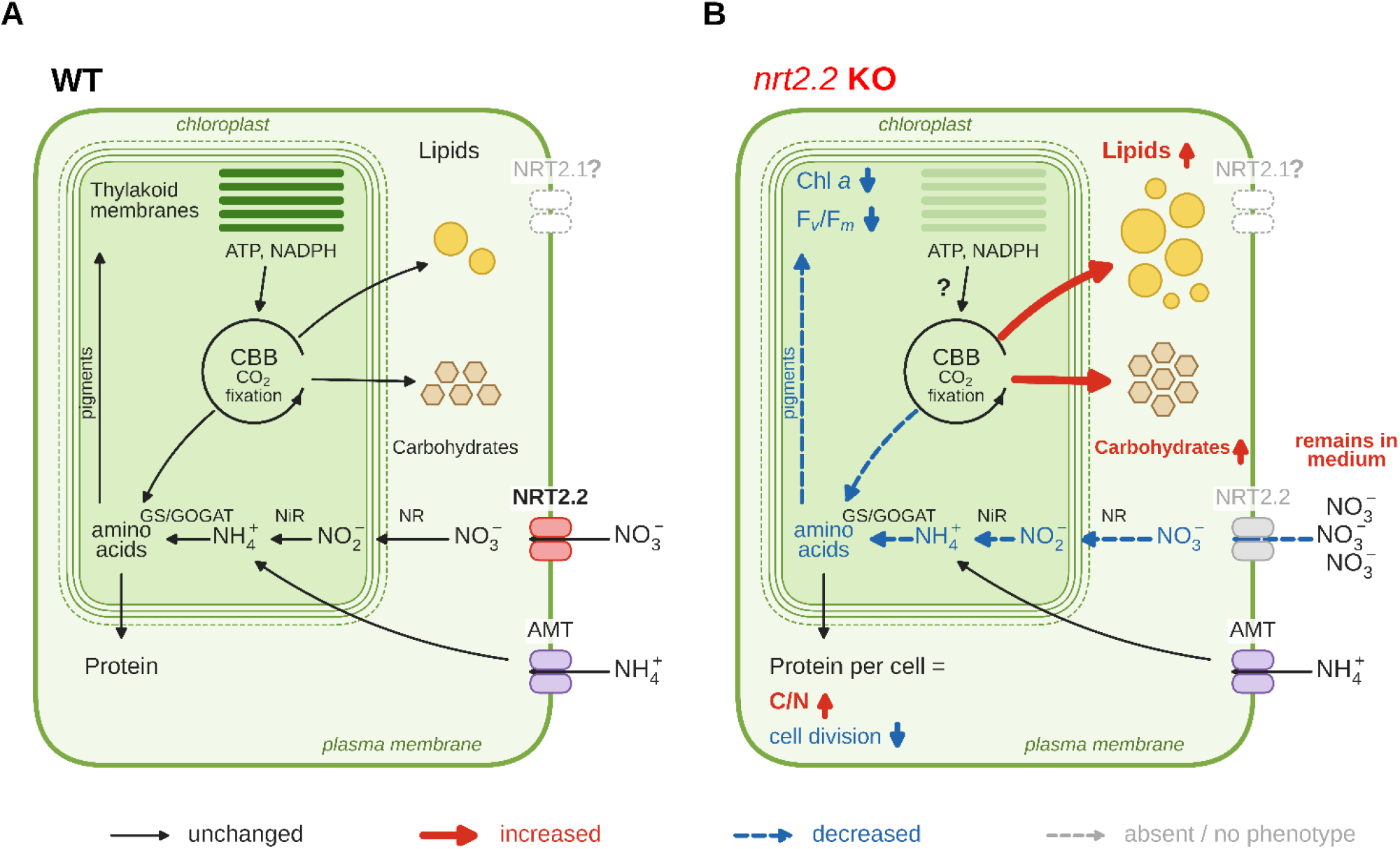
Model of NRT2.2-mediated NO_3_^−^ uptake in *Nannochloropsis oceanica* and consequences of its loss. A) Wild type (WT). NO_3_^−^ enters through the plasma-membrane transporter NRT2.2, is reduced to NO_2_^−^ by nitrate reductase (NR) in the cytosol and to NH_4_^+^ by nitrite reductase (NiR) in the chloroplast, where it is assimilated into amino acids (GS/GOGAT) using carbon skeletons from the Calvin-Benson-Bassham (CBB) cycle. Amino acids sustain protein synthesis. NRT2.1 (grey) is not required for NO_3_^−^ uptake under these conditions. B) *nrt2.2* knock-out (KO). Without NRT2.2, NO_3_^−^ is not taken up and remains in the medium, and cells become N-limited although NO_3_^−^ is available: chlorophyll *a* decreases (paler thylakoids) together with F_v_/F_m_, cell division stops and fixed carbon is diverted to storage lipids and carbohydrates (Fig. 4), increasing the C/N ratio; protein content per cell is unaffected. NH_4_^+^, taken up independently of NRT2.2 (AMT, ammonium transporter), restores growth. Black arrows, unchanged; thick red arrows, increased; dashed blue arrows, decreased; grey, absent or no phenotype. The areas of the lipid droplets (yellow) and carbohydrate symbols (hexagons) are proportional to the FTIR ratios in +NO_3_^-^ (Fig. 4D,E). ATP and NADPH supply to the CBB cycle was not measured (question mark). The outermost chloroplast membrane (dashed) is continuous with the endoplasmic reticulum. ALT TEXT: Model comparing carbon and nitrogen metabolism in wild-type and *nrt2.2* knock-out cells.

### NRT2.2 at the interface between the ocean and the cell

NRT2.2 is the plasma membrane gateway for NO_3_^-^ in *N. oceanica*. From a physiological perspective, these findings place NO_3_^-^ uptake upstream of the coordination between C and N metabolism. By controlling NO_3_^-^ entry, it determines whether fixed C is channelled into growth or diverted into storage, placing it at the core of the metabolic response to N availability (Fig. 6). Because *N. oceanica* relies on a single dominant transporter, its metabolism has little buffering against the loss of this uptake system.

This tight coupling also has ecological implications. In the ocean, NO_3_^-^ is the largest pool of bioavailable N, yet it is depleted for long periods in surface waters and resupplied episodically (Gruber, 2008). Under these conditions, the capacity to acquire NO_3_^-^ can strongly limit primary productivity, and the dependence of *N. oceanica* on a single dominant transporter makes NRT2.2 a strong candidate to determine how this alga uses transient NO_3_^-^ inputs and how it then allocates C between growth and storage, consequently affecting the whole community. Moreover, macromolecular composition determines cell density and, in non-motile phytoplankton, sinking: carbohydrates and proteins increase density, whereas lipids lower it (Liefer et al., 2019; Wu et al., 2025). In nutrient-depleted surface waters, protein and carbohydrate ballast has been proposed to help cells sink toward deeper, nutrient-rich layers (Wu et al., 2025). The concurrent accumulation of carbohydrates and lipids in the *nrt2.2* KO lines may therefore alter buoyancy, thus impairing sinking behaviour. By linking nutrient supply to the allocation of fixed C, NRT2.2 connects the chemistry of seawater to the biochemical composition of the cell, with potential consequences for the C:N stoichiometry of the organic matter produced in the ocean (Liefer et al., 2019).

Together, these results show that NRT2.2 is the gateway through which NO_3_^-^ enters the cell and sets the balance between growth and C storage, providing a mechanistic link between the N supply of the ocean and the metabolism of phytoplankton in a fluctuating environment. Defining how this gateway is regulated and how widely it is shared among lineages will be key to predicting how phytoplankton cope with a variable N supply and, given the biotechnological relevance of *Nannochloropsis*, to redirect C partitioning toward storage compounds such as lipids.

## DATA AVAILABILITY STATEMENT

All data have been deposited in the Research Data UniPD repository under the following accession ID: https://researchdata.cab.unipd.it/id/eprint/1961.

## ACKNOWLEDGEMENTS

The authors wish to thank the Imaging Light Microscopy Facility and the Plant Genome Editing and Phenotyping Facility of the Biology Department of the University of Padova (Italy). AI was used to assist with figure assembly and the scripts used to plot the data without altering the image data, and to revise the English language of the manuscript without changing its scientific content.

## AUTHOR CONTRIBUTIONS

Anna Santin: Conceptualization, Data curation, Formal analysis, Investigation, Methodology, Visualization, Writing - original draft, Writing - review & editing

Laura Verdesca: Formal analysis, Investigation, Methodology, Writing - review & editing Alessandra Bellan: Formal analysis, Investigation, Methodology, Writing - review & editing Miles Minio: Data curation, Formal analysis, Investigation, Methodology, Visualization, Writing - review & editing

Caterina Gerotto: Data curation, Funding acquisition, Investigation, Resources, Supervision, Validation, Writing - review & editing

Tomas Morosinotto: Funding acquisition, Supervision, Writing - review & editing Giorgio Perin: Funding acquisition, Supervision, Writing - review & editing

Alessandro Alboresi: Conceptualization, Data curation, Funding acquisition, Investigation, Project administration, Resources, Supervision, Validation, Writing - review & editing

