## Supplemental Figures for "NRT2.2 is essential for nitrate uptake and carbon-nitrogen balance in *Nannochloropsis oceanica*"

**TITLE**

**AUTHOR NAMES AND AFFILIATIONS**

Anna Santin [1,a], Laura Verdesca [1,b], Alessandra Bellan [1], Miles Minio [2], Caterina Gerotto [2], Tomas Morosinotto [1], Giorgio Perin [1], Alessandro Alboresi [1]

[1] Department of Biology, University of Padova, 35131, Padova, Italy

[2] Department of Life and Environmental Sciences, Marche Polytechnic University, 60131 Ancona, Italy

[a] present address: UMR7141- Photobiology and Physiology of Plastids and Microalgae (P3M), CNRS, Sorbonne University, 75005, Paris, France

[b] present address: Stazione Zoologica Anton Dohrn, Villa Comunale, 80121, Naples, Italy

### SUPPLEMENTARY FIGURES

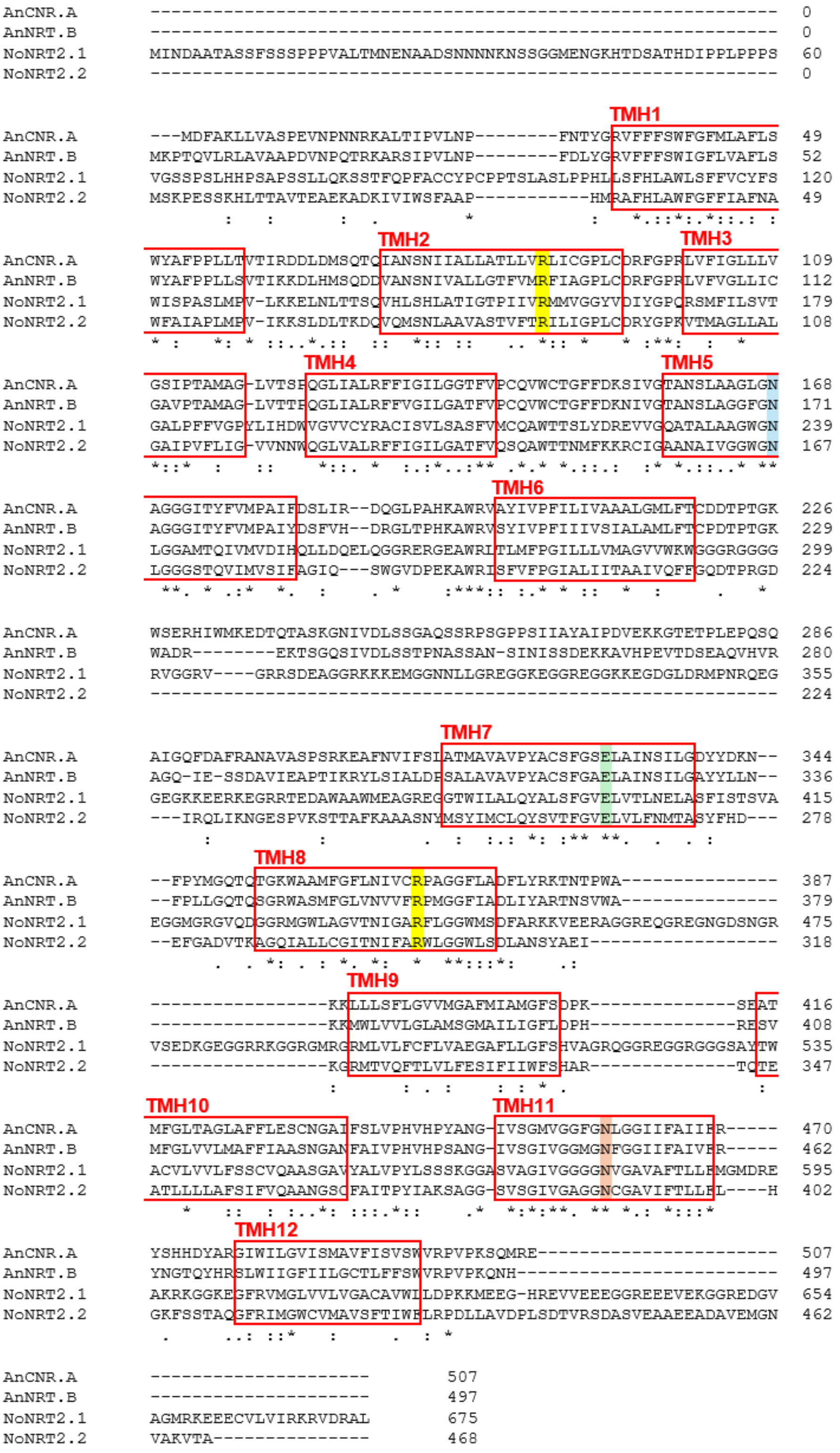


**Figure S1. Structural features inferred from *Nannochloropsis oceanica* NRT2 protein sequences.** Alignment of *N. oceanica* NRT2.1 and NRT2.2 with *Aspergillus nidulans* NrtA (formerly CrnA) and NrtB, showing the 12 transmembrane domains and the key residues putatively involved in substrate binding, as previously identified (Unkles *et al.*, 2004).

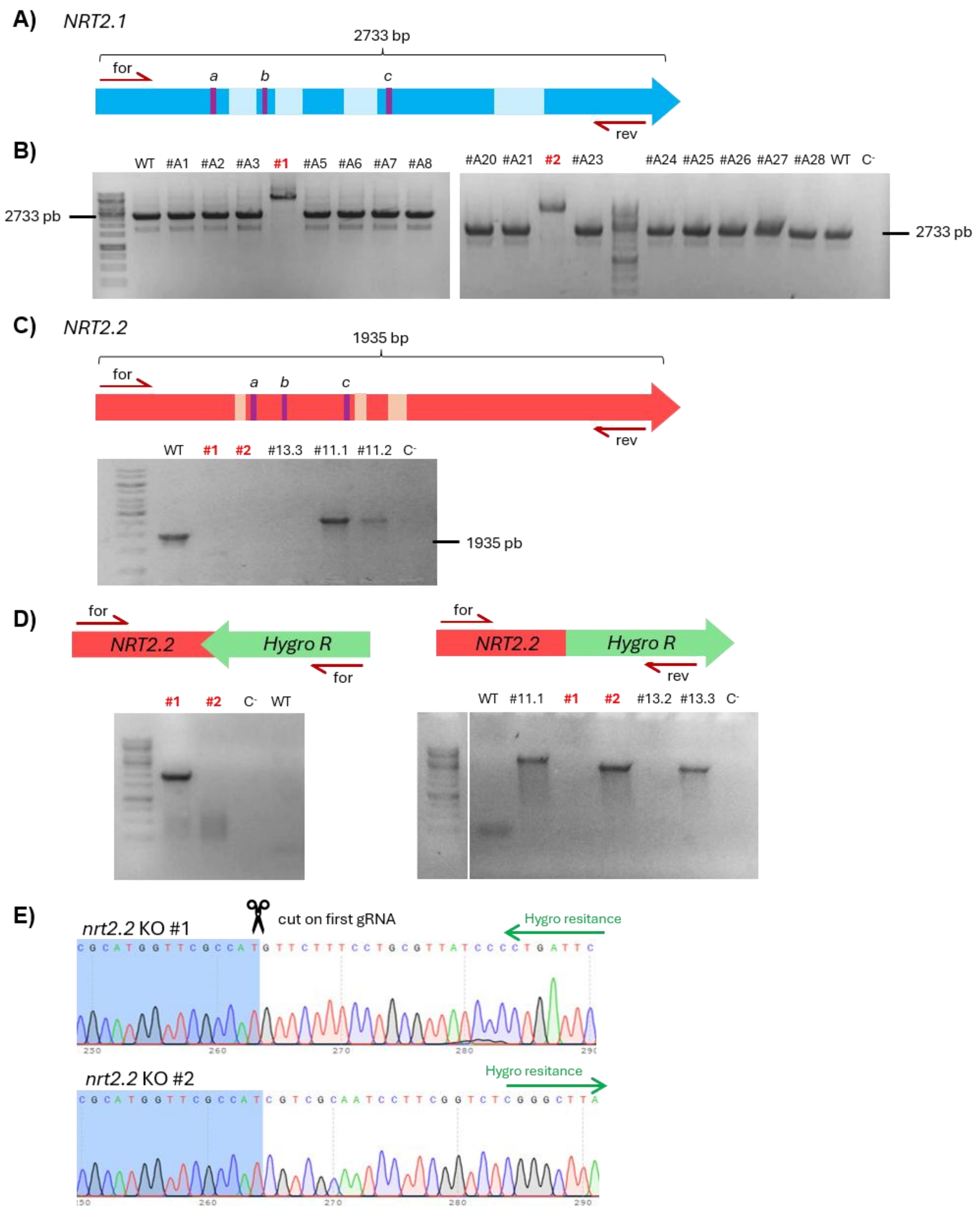


**Figure S2.** ***Nannochloropsis oceanica*** **NRT2 knock-out mutant generation.** A) NRT2.1 gene structure with introns (light blue), gRNA target sites (a-c, purple bars) and primers (for/rev, red arrows). B) Genomic PCR of *nrt2.1* transformants: insertion of the hygromycin resistance cassette gives a longer product; knock-out (KO) lines #1 and #2 (red) were selected. C) NRT2.2 gene structure (introns in light orange) and genomic PCR of *nrt2.2* transformants: no product is amplified in KO lines #1 and #2 (red). D) PCR with one primer on NRT2.2 and one on the cassette, showing insertion in antisense orientation in *nrt2.2* KO #1 (left) and in sense orientation in *nrt2.2* KO #2 (right). E) Sequencing of the insertion sites in *nrt2.2* KO #1 and #2. WT, wild-type parental strain; C-, blank.

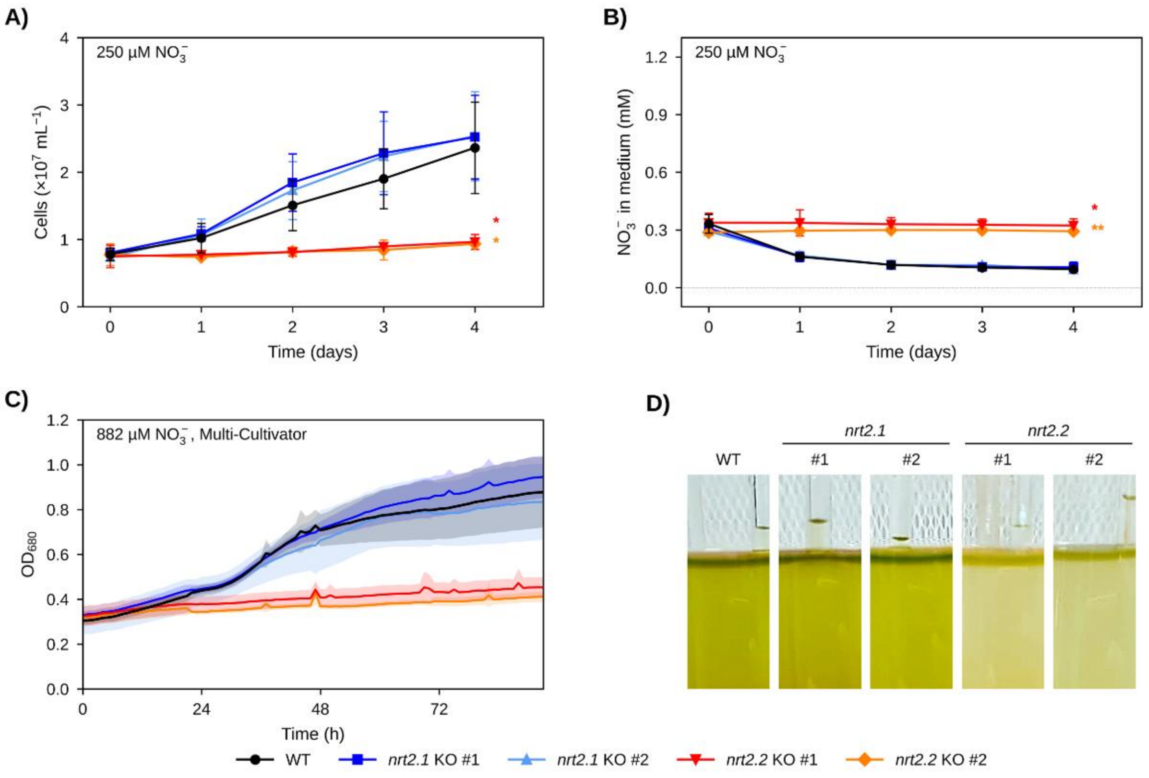


**Figure S3. Growth of *Nannochloropsis oceanica nrt2* knock-out mutants at low $NO_3^-$ and in the Multi-Cultivator system.** A) Cell concentration and B) residual $NO_3^-$ in the medium of wild-type (WT), two *nrt2.1* and two *nrt2.2* knock-out (KO) lines during 4 days in 250 µM $NO_3^-$ in flasks, as in Figure 2 (n = 3 for A, n = 2 for B). **C)** Growth, represented as optical density ($OD_{680}$), monitored every hour in the Multi-Cultivator system under atmospheric $CO_2$ (400 ppm) and continuous white light (80 µmol photons $m^{-2}$ $s^{-1}$) after repletion with 882 µM $NO_3^-$ (mean ± SD, shaded). **D)** Images of the Multi-Cultivator cultures after 4 days of $NO_3^-$ repletion. Asterisks in A and B: unpaired two-tailed Student's t-test of each KO line *vs* WT at day 4; * $p < 0.05$, ** $p < 0.01$.

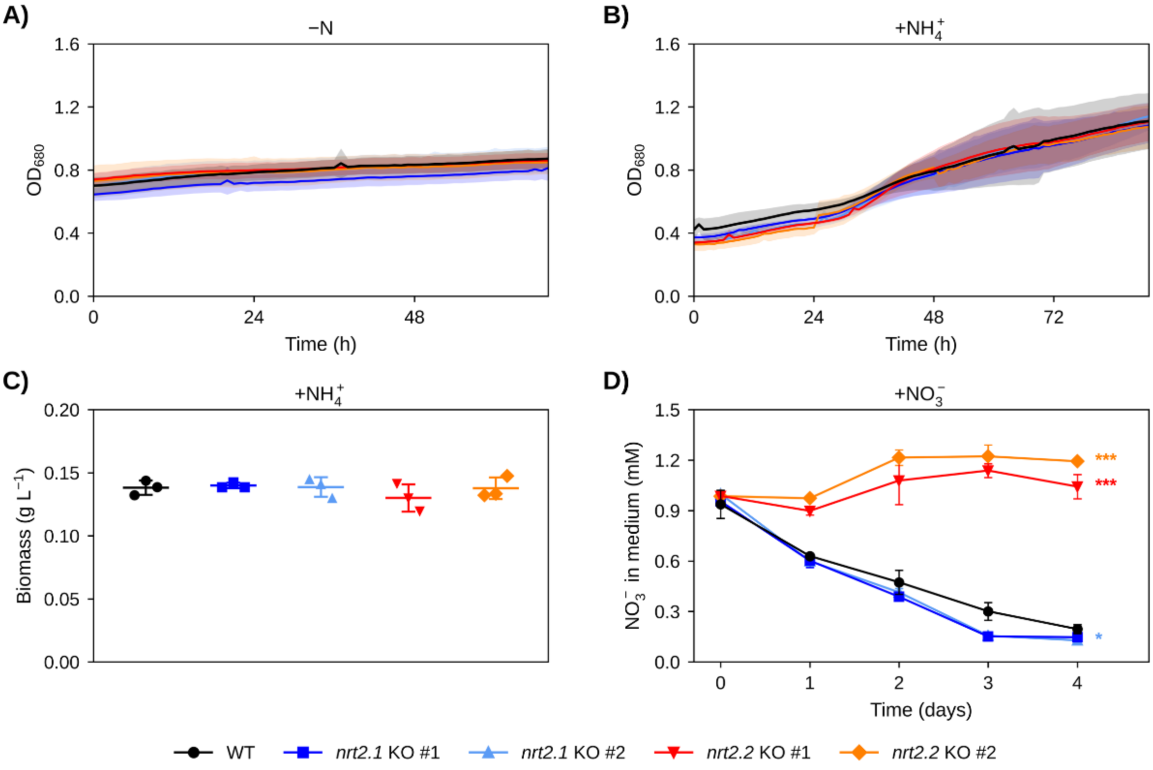


**Figure S4. Growth and $NO_3^-$ uptake of *Nannochloropsis oceanica nrt2* knock-out mutants in the Multi-Cultivator control conditions. A-B)** Growth, represented as optical density ($OD_{680}$) monitored every hour, of wild-type (WT), two *nrt2.1* and two *nrt2.2* knock-out (KO) lines grown in the Multi-Cultivator system under atmospheric $CO_2$ (400 ppm) and continuous white light (80 µmol photons $m^{-2}$ $s^{-1}$) during 72 h of N deprivation **(A)** and after repletion with 882 µM $NH_4^+$ **(B)**. **C)** Biomass production after 4 days of $NH_4^+$ repletion. D) Residual $NO_3^-$ in the medium during 4 days in 882 µM $NO_3^-$. Growth and biomass in +$NO_3^-$ are shown in Fig. S3C and Fig. 3A. Data are mean ± SD of three independent biological replicates (shaded areas in A-B). Statistical significance *vs* WT (Student's t-test; in D at day 4): * $p < 0.05$, ** $p < 0.01$, *** $p < 0.001$.

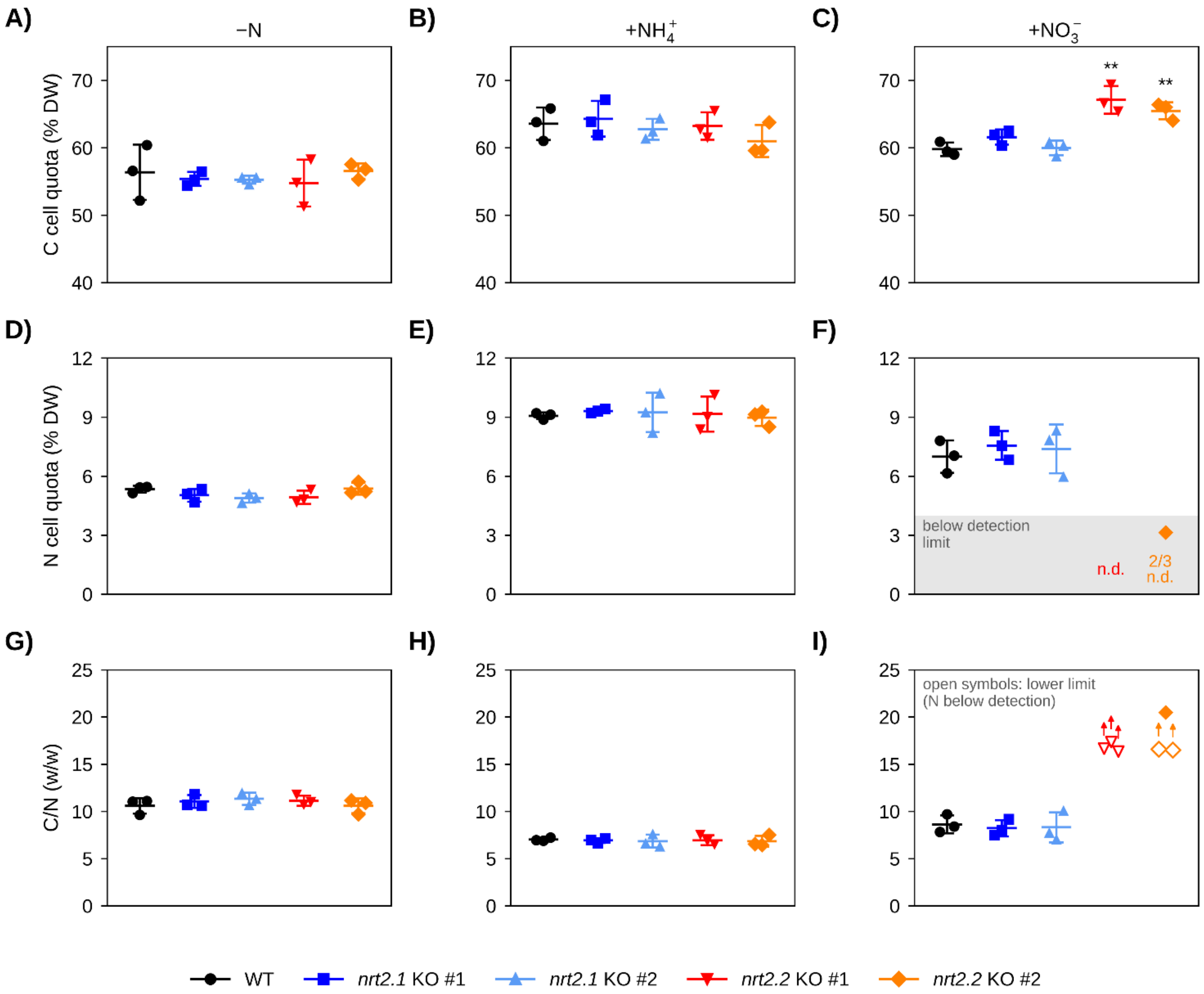


**Figure S5. Intracellular carbon and nitrogen content of *Nannochloropsis oceanica nrt2* knock-out mutants. A-C)** Intracellular carbon (C), **D-F)** intracellular nitrogen (N), as percentage of dry weight, and **G-I)** C/N mass ratio of wild-type (WT), two *nrt2.1* and two *nrt2.2* knock-out (KO) lines grown as in Figure 3, after 72 h of N deprivation (−N) or 4 days of repletion with 882 µM ammonium ($+NH_4^+$) or nitrate ($+NO_3^-$) (A, D, G: −N; B, E, H: $+NH_4^+$; C, F, I: $+NO_3^-$). $+NO_3^-$ data are the same as in Fig. 3B,C. In $+NO_3^-$, N was below the detection limit of the instrument (grey area, 4%) in five of six *nrt2.2* KO samples (n.d.); for these samples C/N is shown as a lower limit (C divided by the detection limit, open symbols). Data are mean ± SD of three independent biological replicates. Statistical significance *vs* WT (Student's t-test): * $p < 0.05$, ** $p < 0.01$, *** $p < 0.001$.

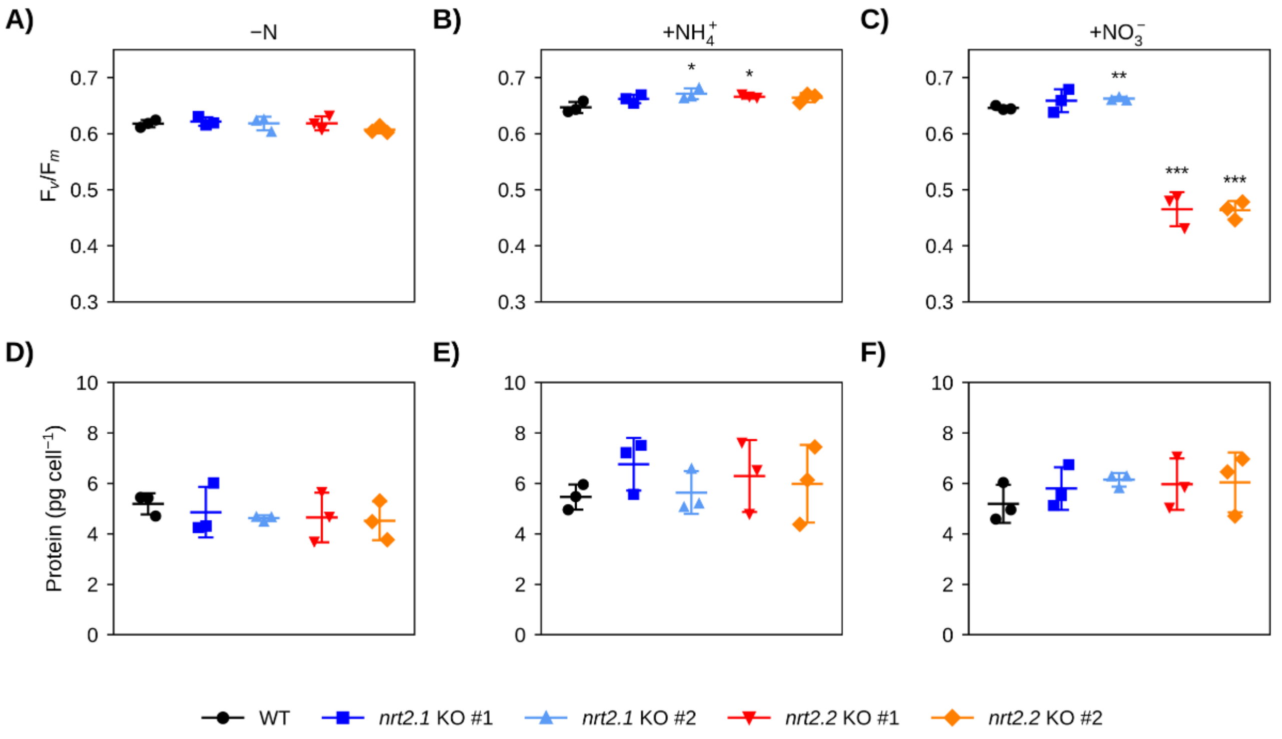


**Figure S6. Photosynthetic efficiency and protein content of *Nannochloropsis oceanica nrt2* knock-out mutants. A-C)** Maximum PSII quantum yield ($F_v/F_m$) and **D-F)** total protein content of wild-type (WT), two *nrt2.1* and two *nrt2.2* knock-out (KO) lines grown as in Figure 3, after 72 h of N deprivation (−N) or 4 days of repletion with 882 µM ammonium ($+NH_4^+$) or nitrate ($+NO_3^-$) (A, D: −N; B, E: $+NH_4^+$; C, F: $+NO_3^-$). $+NO_3^-$ data are the same as in Fig. 3D,E. Data are mean ± SD of three independent biological replicates. Statistical significance *vs* WT (Student's t-test): * $p < 0.05$, ** $p < 0.01$, *** $p < 0.001$.

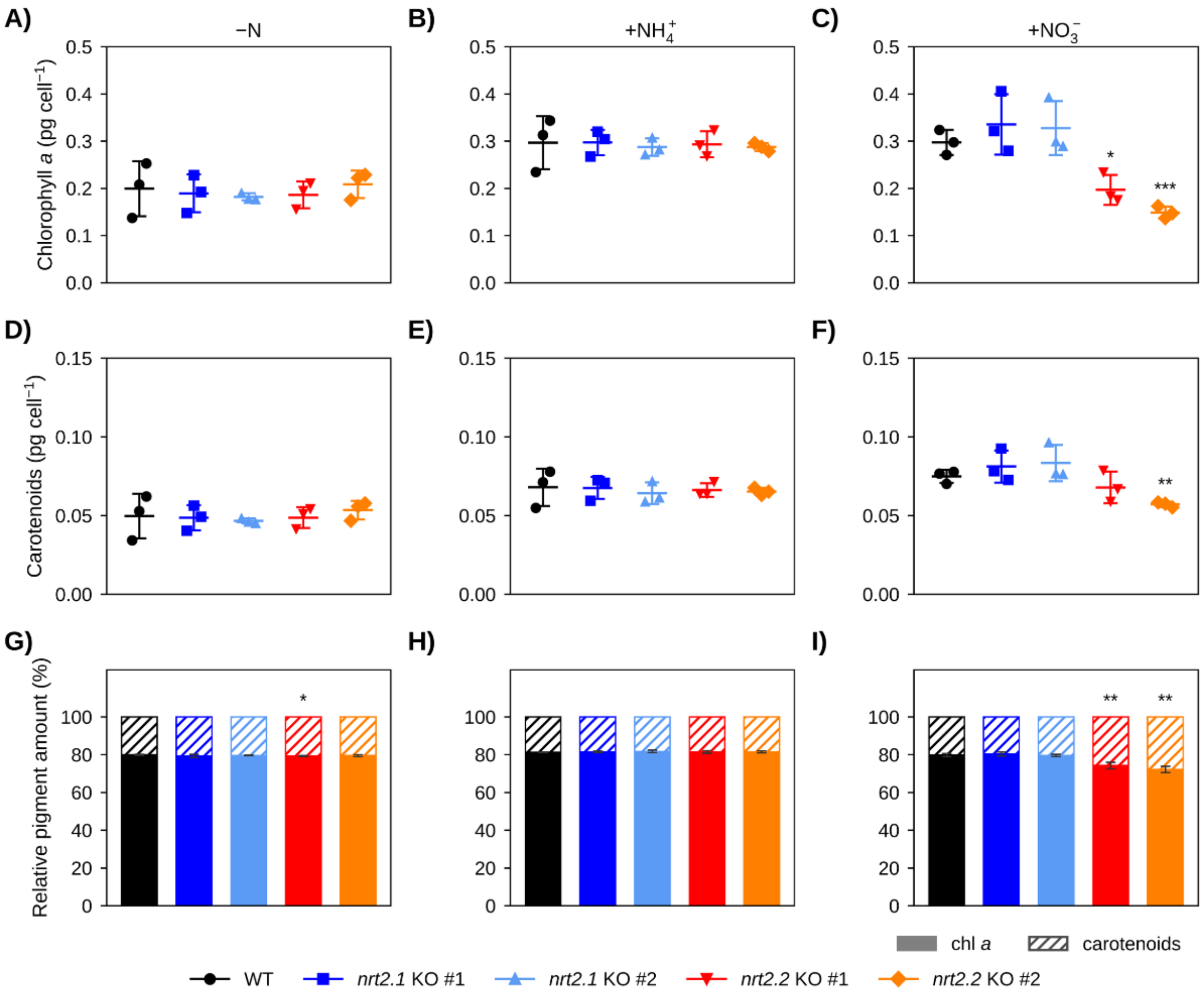


**Figure S7. Pigment content of *Nannochloropsis oceanica nrt2* knock-out mutants. A-C)** Chlorophyll *a* and **D-F)** total carotenoid content per cell, and G-I) relative pigment composition (chlorophyll *a*, filled; carotenoids, hatched) of wild-type (WT), two *nrt2.1* and two *nrt2.2* knock-out (KO) lines grown as in Figure 3, after 72 h of N deprivation (−N) or 4 days of repletion with 882 µM ammonium (+$NH_4^+$) or nitrate (+$NO_3^-$) (A, D, G: −N; B, E, H: +$NH_4^+$; C, F, I: +$NO_3^-$). Data in C and I are the same as in Fig. 3F,G. Data are mean ± SD of three independent biological replicates. Statistical significance *vs* WT (Student's t-test; in G-I on the chlorophyll *a* fraction): * $p < 0.05$, ** $p < 0.01$, *** $p < 0.001$.

### SUPPLEMENTARY TABLES

**Table S1.** Oligonucleotides: crRNAs designed for CRISPR/Cpf1 transformation (with PAM sequence underlined), primers used for NRT2.2-mVenus plasmid construction, and primers used to screen NRT2.2-mVenus and knock-out (KO) lines.

| Category | Name | Sequence |
|---|---|---|
| crRNAs | gNRT2.1_a | 5’- TTTAAGACAGGCATGAGCGACGCTG -3’ (-) |
| | gNRT2.1_b | 5’- TTTGTGGGGCCCTATTTGATCCACG -3’ (+) |
| | gNRT2.1_c | 5’- TTTCCGGGTATCTTGCTGCTGGTCA -3’ (+) |
| | gNRT2.2_a | 5’- TTTATGACGGGCATAAGGGGGGCGA -3’ (-) |
| | gNRT2.2_b | 5’- TTTATAGGCATTTTGGGTGCTACAT -3’ (+) |
| | gNRT2.2_c | 5’- TTTCCCTGGTATTGCCCTCATTATC -3’ (+) |
| Primers for KO screening | NoNRT2.1_cds_for | ATGATAAATGATGCAGCAACAG |
| | NoNRT2.1_cds_rev | CTATAAAGCCCTGTCTACCCTT |
| | NoNRT2.2_cds_for | ATGTCCAAGCCTGAGTCATCCA |
| | NoNRT2.2_cds_rev | TTAGGCGGTGACCTTCGCTACA |
| | NoNRT2.2_cds_for_2 | ATGCCCACGCGCGGCC |
| | Hyg_for | AAGATGGAGTGGATGGAGGAGGA |
| | Hyg_rev | TCTAGTGGATCAGCTTGCATGC |
| Primers for OE plasmid construction | pNoLdsp_yfp_his_backbone_fwd | AGTAGATGCCGACCGGAG |
| | pNoLdsp_yfp_his_backbone_rev | ATCTTATTGCCCCCCGGG |
| | pNoLdsp_yfp_his_NRT2.2_cds_fwd | GCCCCGGGGGGCAATAAGATATGTCCAAGCCTGAGTCATC |
| | pNoLdsp_yfp_his_NRT2.2_cds_rev | GGCGGTGACCTTCGCTAC |
| | pNoLdsp_yfp_his_Tags_fwd | ATGTAGCGAAGGTCACCGCCGGTGGTACCATGGTGAGC |
| | pNoLdsp_yfp_his_Tags_rev | GACTCCGGTCGGCATCTACTGACGTCTCAGTGATGGTG |
| Primers for OE screening | Noc_LSDP_prom_scrn_for | GTGTGCCTGGTTGTTCCGCCA |
| | HygR_cass_term_scrn_rev | GCTCAACACATGAGCGAAACCC |
| | YFP_start_rev | CGGACTTGAAGAAGTCGTGCTGC |
